# Network analysis reveals a major role for 14q32 cluster miRNAs in determining transcriptional differences between IGHV-mutated and unmutated CLL

**DOI:** 10.1101/2022.04.26.487675

**Authors:** Dean Bryant, Lindsay Smith, Karly Rai Rogers-Broadway, Laura Karydis, Jeongmin Woo, Matthew D Blunt, Francesco Forconi, Freda K Stevenson, Christopher Goodnow, Amanda Russell, Peter Humberg, Graham Packham, Andrew J Steele, Jonathan C Strefford

## Abstract

Tumour cells from patients with chronic lymphocytic leukaemia (CLL) can express unmutated (U-CLL) or mutated (M-CLL) immunoglobulin heavy chain (IGHV) genes with differing clinical behaviours, variable B cell receptor (BCR) signalling capacity and distinct transcriptional profiles. As it remains unclear to what extent these differences reflect the tumour cells’ innate pre/post germinal centre origin or their BCR signalling competence, we applied RNA sequencing, small RNA sequencing and DNA methylation array analysis to 38 CLL cases categorised into three groups by IGHV mutational status and BCR signalling capacity. We identified 492 mRNAs and 38 miRNAs differentially expressed between U-CLL and M-CLL, but only 9 mRNAs and 0 miRNAs associated with BCR competence within M-CLL. A significant proportion of the IGHV-associated miRNAs derived from chr14q32 clusters (14/38 (37%)), where all miRNAs were co-expressed with the *MEG3* lncRNA, as part of the DLK1-DIO3 genomic imprinted region, a locus of known importance in the pathogenesis of other human tumours. Integrative *in silico* analysis of miRNA/mRNA data revealed pronounced regulatory potential for the 14q32 miRNAs, potentially accounting for up to 25% of the IGHV-related transcriptome signature. GAB1, a positive regulator of BCR signalling, was predicted to be regulated by five 14q32 miRNAs and we confirmed that two of these (miR-409-3p and miR-411-3p) significantly repressed activity of the *GAB1* 3’UTR. Our analysis demonstrates a potential key role of the 14q32 miRNA locus in the regulation of CLL-related gene regulation.

## Introduction

Chronic lymphocytic leukaemia (CLL) is a heterogeneous neoplasm of mature B-cell origin. Early genetic studies of the immunoglobulin heavy-chain variable region (IGHV), that encodes part of the B-cell receptor (BCR), identified two disease subtypes based on the degree of somatic hypermutation (SH) of the IGHV locus whereby CLL with mutated IGHV genes (M-CLL) and unmutated IGHV genes (U-CLL) typically have a favourable and poorer prognosis, respectively^1,2^. Subsequent gene expression, DNA methylation and chromatin accessibility studies support a distinct cell of origin for these two CLL subtypes, with M-CLL and U-CLL exhibiting the expression profile of post-germinal centre memory cell, and pre-germinal centre B-cell, respectively^3–6^. In addition, DNA methylation profiling further identified a third minor subgroup exhibiting an intermediate methylome between M-CLL and U-CLL, with distinct immunogenetic and genomic features and intermediate clinical outcome^7–9^.

The B-cell receptor (BCR), as evidenced by the efficacy of BTK inhibition, is of critical importance to the progression of CLL^10^. CLL cells typically express immunoglobulin D (IgD) and M (IgM) on their cell surface, although at reduced levels compared to normal B cells and CLL is hypothesised to undergo BCR engagement by autoantigen^11^. In fact, within this low range, tumour cells from CLL patients exhibit considerable variation in both the level of surface IgM (sIgM), and in their capacity to signal via anti-IgM engagement. Whilst U-CLL typically exhibit higher sIgM levels and signalling capacity than M-CLL, there is overlap and higher functional levels of sIgM have been associated with patient survival independent of IGHV status; signalling competent U-CLL had a median survival of 32 months, signalling competent M-CLL had a median survival of 81 months and signalling deficient M-CLL had a median survival of 183 months^12^. The outcome of BCR signalling can range from B cell activation to anergy, with perhaps more of the latter in M-CLL^13^. However, how the differential responses to anti-IgM stimulation of the BCR operate *in vivo* is not yet fully understood.

MicroRNAs (miRNAs) are a class of short, ^~^22nt long non-coding RNAs with important roles in the regulation of gene expression. Mapping miRNA *loci* at the genome level shows that miRNAs are often intragenic, located in introns, occur within clusters of co-regulated miRNAs and are expressed as a consequence of host gene transcription^14^. Most miRNAs are transcribed from DNA into primary miRNA prior to further processing in to precursor and mature species (reviewed in O’Brien *et al*. 2018^15^). miRNAs have been shown to influence a network of transcription factors that regulate B-cell differentiation, initially demonstrated by the conditional knockdown of *AGO2* and Dicer, that resulted in impaired B-cell maturation, impaired germinal centre formation, aberrant apoptosis and enhanced BCR signalling^16–18^. Analysis of 13q14 deletion in CLL was the first reported link between somatic genomic lesions and deregulation of miRNAs, specifically miR-15a and miR-16-1, which are negative regulators of the antiapoptotic proteins Bcl-2 and Mcl-1^19,20^. miR-34b/c, miR-21, miR-29, miR-125b, miR-181b, miR-17/92, miR-150, and miR-155 family miRNAs are also suggested to be biologically and clinically important in CLL, are differentially expressed compared to the normal B-cells^21^ and can be utilised to risk-stratify CLL patients into prognostically relevant subtypes^22^. Several miRNAs have been causally linked to CLL pathogenesis, including miR-15a/16-1, miR-29b/miR-181b, miR-181b and miR-34a that interact with the anti-apoptotic and cell cycle control genes *BCL2, TCL1, MCL1* and *TP53* respectively^23–26^. miR-150 and miR-155 are the most abundantly expressed miRNAs in CLL, where they regulate BCR signalling by targeting *FOXP1* and *GAB1* mRNAs and via downmodulation of the BCR phosphatase *SHIP1*, respectively^27,28^. However, there remains uncertainty as to what degree the abnormal miRNAs reflect the cause or consequence of disrupted BCR signalling.

Whilst considerable research focus has described the biological basis underpinning the clinical heterogeneity observed in M-CLL and U-CLL, it remains unclear to what extent these differing biological features reflect the tumour cells innate pre/post-germinal centre origin or their acquired BCR signalling competence. Consequently, we performed a detailed multi-omics analysis of CLL samples differentiated by IGHV mutational status and BCR signalling capacity. By employing genome-wide miRNA and mRNA sequencing, with integration of matched DNA methylation, copy number analysis and sophisticated data integration approaches, we identify differential expression of a cluster of miRNAs within the *DLK1-DIO3* imprinted locus at 14q32, that associated with IGHV mutation status. Furthermore, we demonstrate a broad impact for the 14q32 miRNAs on the CLL transcriptome and identify the BCR regulating gene, *GAB1*, as a potential target of multiple 14q32 miRNAs.

## Methods

### Patient cohorts and sample characteristics

An overview of the cohort and methods applied in this study are depicted in Figure 1A (**Fig 1A**). Thirty-eight basal tumour samples, obtained from CLL patients diagnosed using the iwCLL guidelines^29^ and managed at Southampton General Hospital (Southampton, UK), were selected based on IGHV mutational status using established cut-offs^1^ (**Fig S1A**), and BCR signalling capacity with >10%/≤10% anti-IgM-induced Ca^2+^ mobilisation thresholds (**Fig S1B**) determined as described previously^12^. Informed consent was obtained in accordance with the declaration of Helsinki and the study was approved by our regional research ethics committee. We focused on high purity (mean 89.5%, range 77%-99%) (**Fig S1C**), but non-purified CLL cells for the initial transcriptome and miRNA sequencing and used purified CLL cells (Miltenyi B-CLL Isolation kit) for DNA methylation analysis and confirmatory miRNA sequencing. CLL cases were divided into three subgroups of BCR-signalling competent U-CLL (U-CLL-S, n=13) and M-CLL (M-CLL-S, n=13) and BCR-signalling deficient M-CLL (M-CLL-NS, n=12). Due to limitations in sample material, GAB1 protein immunoblotting was performed on a subset of our main cohort comprising U-CLL-S (n=5), M-CLL-S (n=6) and M-CLL-NS (n=4), and additional CLL samples with BCR signalling and IGHV data available, including U-CLL-S (n=8) and M-CLL-S (n=3). Detailed methods presented in **Supplemental Methods**.

**Figure 1.**
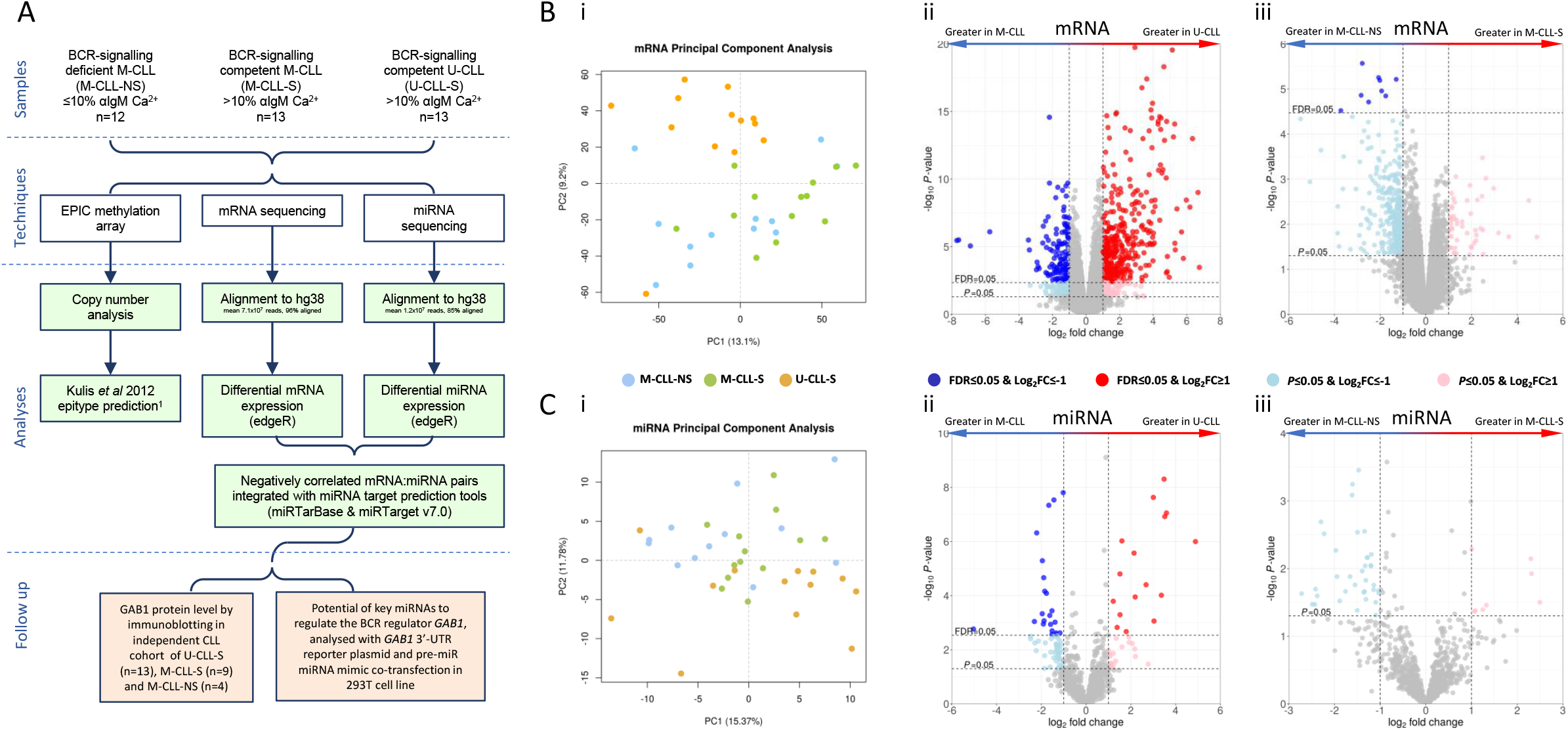
Project outline, PCA plots and volcano plots for pairwise differential expression comparisons. (A) Consort diagram showing the order of investigation of the mRNA, miRNA and DNA methylation profiles of CLL. (B) i) PCA plots using mRNA data and volcano plots for mRNA pairwise comparisons of ii) U-CLL vs. M-CLL and (iii) M-CLL-S vs. M-CLL-NS. (C) i) PCA plots using miRNA data and volcano plots for miRNA pairwise comparisons of ii) U-CLL vs. M-CLL and iii) M-CLL-S vs. M-CLL-NS. Points represent a mRNA or miRNA. Dark blue/red are differentially expressed at FDR≤0.05 and Log_2_FC ≥ +/−1 after correcting for multiple testing (BH), light blue/pink are significant at *P*≤0.05 and Log_2_FC ≥ +/−1.

### mRNA and miRNA sequencing

mRNA libraries were prepared using TruSeq RNA Library Prep kit (Illumina, Hayward, CA, USA) and sequenced using an Illumina HiSeq2500 and HiSeq4000. Raw data was quality checked using FastQC (Babraham Bioinformatics, Cambridge, UK) and aligned to the hg38 reference genome using HISAT2^30^. Read counts per gene were calculated using HTseq-count^31^ against Ensembl GRCh38 v94^32^.

miRNA libraries were prepared using Illumina TruSeq small RNA library kits (Illumina, Hayward, CA, USA) sequenced using an Illumina HiSeq 4000 and an Illumina HiSeq 2500 (Illumina, Hayward, CA, USA). Subsequent, confirmatory miRNA sequencing was performed to preclude any potential contribution of contaminating T-cell/monocytes to our findings, processed as before, but from purified CLL cells (Miltenyi B-CLL Isolation kit) and sequenced on an Illumina HiSeq 4000 (Illumina, Hayward, CA, USA). miRNA data in the form of fastq were quality checked using FastQC, then aligned to the hg38 reference genome using BWA v0.7.12^33^. miRNA read counts were calculated using HTseq-Count against miRbase v21^34^.

### EPIC DNA methylation array

DNA methylation was assessed on purified CLL tumour cells using EPIC DNA methylation arrays (Illumina, Hayward, CA, USA). Data was analysed using RnBeads v2.93^35^ from raw intensity data through to differential methylation analysis and output beta/M values. Following removal of SNP enriched probes (17,371) and unreliable probes using the GreedyCut method (17,302), 832,222 probes were used for the analysis (**Fig S2A**). Conumee^36^ was used to produce copy number profiles. Methylation epitypes were determined as naïve like (n-CLL), memory like (m-CLL) or intermediate (i-CLL) as described by Kulis *et al* ^4^. All other analysis were performed in R v.3.6.1^37^ using custom code.

### Immunoblotting

Immunoblotting was performed as previously described^38^ using rabbit anti-GAB (Cell Signaling Technologies, Danvers, MA, USA) or mouse anti-Hsc70 (Santa Cruz Biotechnology Inc, Dallas, Texas, USA) primary antibodies and horseradish peroxidase-conjugated anti-rabbit/anti-mouse secondary antibodies (Dako, Santa Clara, CA, USA). Representative immunoblots are shown in supplementary Figure 3 (**Fig S3**).

### miRNA transfections

miRNA activity was analysed by co-transfecting 293FT cells (ThermoFisher, Leicestershire, UK) with Lipofectamine 2000 (ThermoFisher, Leicestershire, UK) and a human *GAB1* 3′-UTR reporter plasmid (containing 3770 bp immediately downstream of the end of the *GAB1* ORF cloned into pMirTarget, Origene), a control Renilla luciferase plasmid (Promega) and pre-miR miRNA mimics or pre-miR control 1 (all ThermoFisher, Leicestershire, UK). Luciferase activity was quantified at 24 hrs using the dual-glo luciferase assay system (Promega, Southampton, UK), normalised using Renilla luciferase values from the same well and normalised values for control transfected cells (no pre-miR) were set to 1.0.

### Data analysis

Both mRNA and miRNA data were filtered to remove low expression features (features with ≥1 cpm in ≥3 (mRNA) or ≥2 (miRNA) samples were retained) (**Fig S2B and S2C**). Differential expression amongst CLL subgroups was assessed using EdgeR v3.32.1^39^ and custom R code.

miRNA:mRNA interaction analysis was performed using the R package miRComb^40^ but heavily modified with custom R code and additional miRNA targets databases (including miRTarBase v7.0, IPA expert validation, TargetScan v7.0, miRSVR, miRDB v5.0 and miRRecords^41–45^). Interaction maps were produced using CytoScape v3^46^ to show negatively correlated miRNA:mRNA pairs present in at least one database of miRNA targets. Ingenuity pathway analysis (http://www.ingenuity.com/index.html) and DAVID v8^47^ were used to further interrogate predicted targets. Statistical analysis of miRNA:mRNA interaction counts was performed by quantifying miRNA:mRNA interactions in TargetScan v7.0 and miRDB v5.0 for miRNAs/mRNAs of interest compared to 50,000 cycles of size matched, randomly selected miRNAs/mRNAs using a 1-way student’s t-test. All analyses were performed in R v.3.6.1 using custom code. Data available on request from the authors.

## Results

### IGHV mutational status drive gene expression more strongly than BCR signalling capacity

We performed mRNA sequencing and achieved a mean of 7.13×10^7^ reads (range 4.37×10^7^-1.21×10^8^) per sample, of which 96.0% (range 92.4-98.8%) and 55.6% (range 42.2-64.2%) mapped to hg38 and GENCODE gene annotations, respectively (**Fig 1A**). 15,529 genes passed filtering criteria (**Fig S2B**). Exploratory PCA analysis was able to delineate U-CLL-S, M-CLL-S and M-CLL-NS into three broadly distinct groups, with good separation based on IGHV mutational status (**Fig 1Bi**).

Initially, we focused on the gene expression signature that differentiated M-CLL (regardless of sIgM signalling capacity) from U-CLL and we identified 371 upregulated and 127 downregulated differentially expressed genes (DEGs) at FDR<0.05 (likelihood ratio test) and with a Log_2_FC≥+/-1 (**Fig 1Bii**). Whilst our data showed a high-level of concordance with previously published datasets^48–50^ (**Fig S4A**), confirming 71/114 of the DEGs previously published (**Fig S4B**), we also were able to identify a significant panel of novel DEGs, suggesting greater resolution in our study.

Using KEGG pathway membership, amongst the upregulated DEGs (FDR ≤0.05, Log_2_FC ≥1) in U-CLL we found genes involved in RAS (*GNB4, GAB1, RASAL1, ANGPT2, FGFR1, IGFR1, INSR, KSR2, PLD1, ZAP70, PRKCA*), Wnt (*LRP5, VANGL1, VANGL2, WNT2B, WNT5B, WNT9A, FZD1, PRKCA*), and PI3K-Akt signalling (*GNB4, TCL1A, ANGPT2, CHAD, FGFR1, IGF1R, INSR, ITGB4, ITGB5, PRKCA, PPP2R3B, TNXB*) as well as other genes coding for proteins known to be over-expressed in U-CLL (*ZAP70, LPL*). Downregulated DEGs (FDR≤0.05, Log_2_FC≤−1) included those involved in MAPK (*DUSP2, DUSP8, NR4A1, TNF, MYC*), BCR (*CD86, CD40LG, EGR1, EGR2, EGR3, EGR4*), TLR (*TLR1*) and TNF signalling (*TNF*, *TNFRSF18, TNFRSF9*), as well as several Interleukins (*IL10*, *IL6*). Using the 498 DEGs in IPA and DAVID, we identified upregulated genes involved in Wnt signalling, down regulation of TNF and PTEN signalling genes, and increased expression of genes associated with cellular migration in U-CLL.

To probe the gene expression signature specifically associated with BCR signalling capacity, we compared M-CLL-NS with M-CLL-S, whilst excluding U-CLL. With thresholds at FDR≤0.05 (likelihood ratio test) and with a Log_2_FC≥+/−1, the signature included 9 downregulated (*ITPRIPL2, TNFRSF9, CALHM2, KIR3DL2, MAF, IL1A, CSF1, FNBP1L* and *NT5E*) and 0 upregulated DEGs in M-CLL-S (**Fig 1Biii**). For the 9 DEGs associated with BCR signalling capacity, expression was not significantly associated with a single pathway, suggesting a limited impact of signalling capacity on the transcriptome in M-CLL. Whilst we anticipated detecting fewer DEGs between M-CLL-S and M-CLL-NS due to the reduced sample size, statistical power alone is unlikely to explain the pronounced difference compared to U-CLL vs. M-CLL.

### miRNA expression strongly associates with IGHV mutational status

Next, we quantified miRNA abundance in the same CLL cohort, sequencing a mean of 1.15×10^7^ reads (range 7.02×10^6^-1.91×10^7^) per library, of which 84.6% (range 74.1-92.0%) mapped to hg38 and 20.6% (range 7.4-40.2%) of mapped with high confidence to a mature miRNA annotated in miRBase 21. Of the 2587 mature miRNAs annotated in miRBase 21; 966, 699 and 922 miRNAs were not detectable in any sample, filtered out due to low read counts (defined as <1 counts per million (cpm) in <2 samples) and taken forward for analysis, respectively (**Fig S2C**). Of our 25 most highly expressed miRNAs, 21/25 were observed in the top 50 most highly expressed miRNAs in a published CLL miRNA dataset^51^ and included key miRNA consistently reported as overexpressed in CLL compared to normal B cells (i.e. miR-150-5p, miR-155-5p, miR-146b-5p, miR-21-5p and miR-29a-3p^22,52^, clearly demonstrating the ability of our approach to validate and extend on established findings.

As we previously observed at the mRNA level, exploratory PCA of our miRNA data showed stronger separation of samples by IGHV mutation status rather than BCR signalling capacity in M-CLL (**Fig 1Ci**). Differentially expressed miRNA (DEM) analysis for IGHV mutation status identified 38 DEMs (16 downregulated and 22 upregulated) at FDR≤0.05 and Log_2_FC≥+/-1 (**Fig 1Cii**), but 0 DEMs for BCR signalling capacity (**Fig 1Ciii**). Comparison of our U-CLL vs. M-CLL miRNA expression signature with the key miRNAs characterised in CLL showed that 12 of the 17 miRNAs associated with IGHV status^53^ behaved similarly in our study, and included the key CLL miRNAs miR-150-5p and miR-146a-5p, down- and up-regulated in U-CLL, respectively^22,54^ (**Fig S4C**). Further characterisation of the 38 DEMs showed several with established roles in cancer. Of the miRNAs downregulated in U-CLL, miR-146b-3p and miR-146b-5p are downregulated in pancreatic cancer and aggressive lung cancer, respectively^55,56^. Of those upregulated in U-CLL, miR-944, miR-138-5p and miR-338-3p are upregulated in advanced cervical cancer^57^, over-expressed in bladder cancer^58^ and thought to have tumour suppressor potential in lung cancer^59^, respectively.

### 14q32 miRNA clusters is differentially expressed in IGHV subgroups

As miRNAs are often located in clusters, we positioned the IGHV-associated DEMs on to the genome and noted that 37% (14/38) mapped to two miRNA clusters in 14q32.2-q32.31, in a ^~^245 kbp locus between the *DLK1* and *DIO3* genes (hg38 coordinates: chr14:100,825,000-101,070,000), approximately 4.5 Mbp upstream of the *IGH* loci (**Fig 2A**). This locus is commonly referred to as the *DLK1-DIO3* imprinted region as it is imprinted during embryonic development, and the lncRNA *MEG3* and downstream 14q32 miRNA clusters are transcribed from the maternal allele^60^. As the 14q32 miRNA clusters contain many miRNAs, only some of which were called as DEMs and miRNA clusters are often co-transcribed, we sought to understand expression of the 14q32 miRNA clusters in their entirety. The 14q32 clusters (*P*=0.0004, Wilcoxon rank sum exact test) and 92.5% of the constituent miRNAs (49/53) were under-expressed in U-CLL compared to M-CLL (**Fig 2B**); the 4/53 miRNAs that failed to show this correlation exhibited low expression levels and/or small Log_2_FC changes. We also observed an association with signalling capacity, where the majority of 14q32 miRNAs were downregulated in M-CLL-S compared to M-CLL-NS, albeit these differences did not reach significance at FDR≤0.05 (**Fig 2Cii**). Taken together, we observed high (mean Log_2_cpm=2.37), intermediate (mean Log_2_cpm=2.04) and low expression (mean Log_2_cpm=1.75) levels of the 14q32 miRNAs in the M-CLL-NS, M-CLL-S and U-CLL-S subgroups, respectively (**Fig 2Ci**), and this was true for miRNAs across the locus (**Fig 2D**).

**Figure 2.**
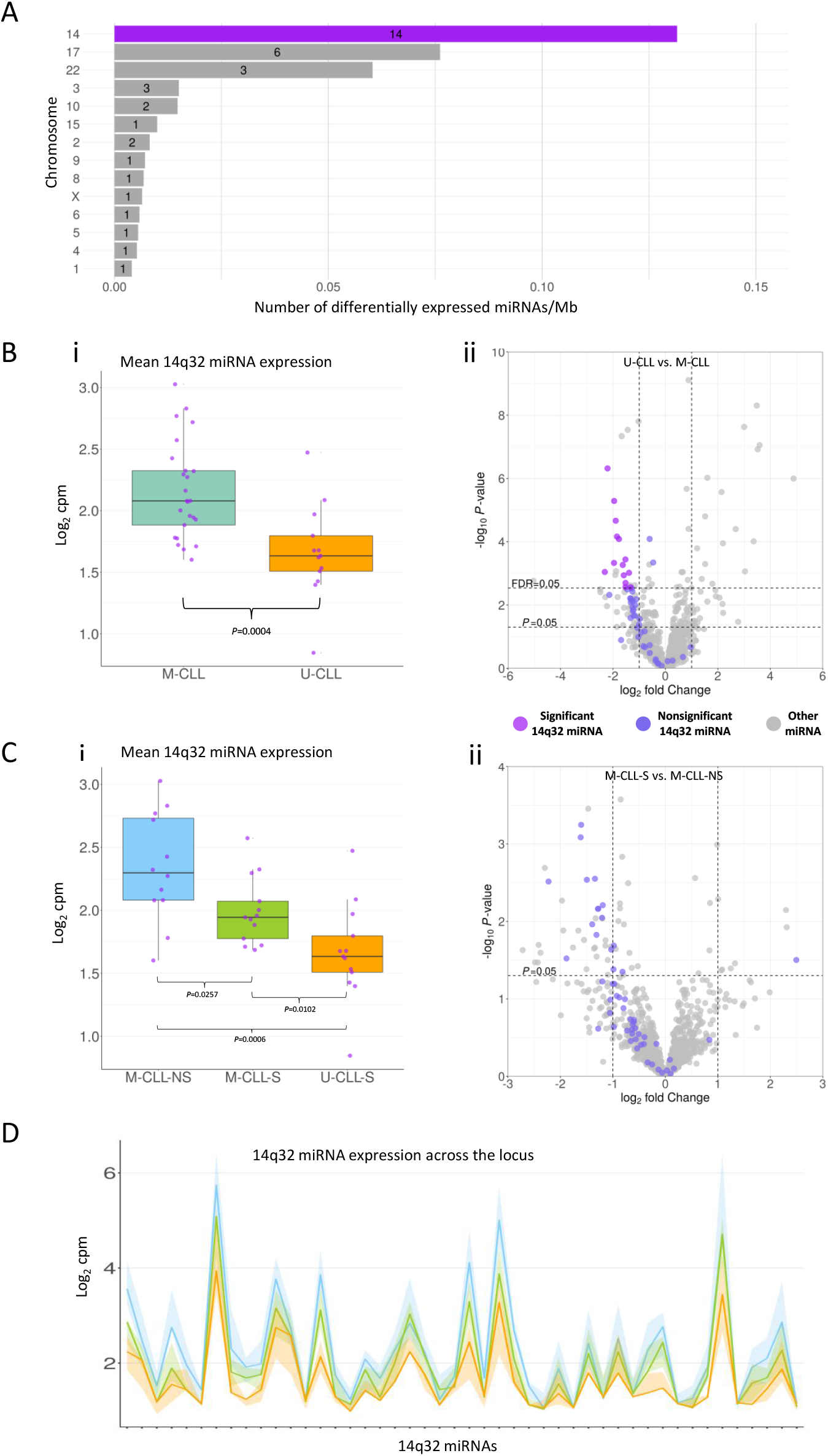
Identification of and expression of the 14q32 miRNA cluster in CLL. (A) Over a third of the differentially expressed U-CLL vs M-CLL miRNAs were located in 14q32 miRNA clusters. Number in bar indicates number of DE miRNAs per chromosome. (B) i) Mean miRNA expression of the 14q32 locus is lower in U-CLL than in M-CLL. ii) The majority of the 14q32 miRNAs (purple points) were downregulated in U-CLL compared to M-CLL. (C) i) Mean miRNA expression of the 14q32 locus is lower in M-CLL-S than in M-CLL-NS and shown to decrease from M-CLL-NS to M-CLL-S to U-CLL-S. ii) The majority of the 14q32 miRNAs (purple points) were downregulated in M-CLL-S compared to M-CLL-NS. (D) Ribbon plot showing the expression of each miRNA in the 14q32 locus ordered by genomic loci from 5’-3’. Dark lines indicate the group median for each miRNA, connection between points is for clarity and no relationship between adjacent miRNAs is inferred. Coloured ribbons indicate the interquartile range for each group. miRNAs plotted include only those expressed at sufficient level to be reliably detected (see methods).

Next, we confirmed this 14q32 miRNA expression distribution using unsupervised k-means clustering. Using this approach, we identified 3 clusters of expression (**Fig S5A**), thereby demonstrating that miRNA expression of the 14q32 clusters could divide patients into three distinct groups that closely aligned with our 3 CLL groups. The high (k-means group 1), intermediate (k-means group 2) and low expression (k-means group 3) clusters included 6/8 M-CLL-NS, 8/14 M-CLL-S, and 10/16 U-CLL-S, respectively. Notably, the outlying U-CLL-S sample in k-means group 1 had 98.1% IGHV homology to germline, just above the 98% threshold at which M-CLL was defined (**Fig S1A**).

Epitype assignment based on DNA methylation as reported by Kulis et al 2012^4^ showed that the majority of U-CLL and M-CLL were assigned to the n-CLL (10/13 U-CLL-S) and m-CLL (10/13 M-CLL-S and 11/12 M-CLL-NS) epitypes respectively, with a small number from each CLL subgroup assigned to the i-CLL epitype (3/13 U-CLL-S, 3/13 M-CLL-S and 1/12 M-CLL-NS). When we overlaid the epitype classes on to our 14q32 miRNA expression clusters, we found that k-means group 1 with the highest and k-means groups 3 with the lowest 14q32 miRNA expression were overrepresented by m-CLL (7/8) and n-CLL (8/16) cases, respectively (**Fig S5A and S5B**).

A striking observation that resulted from this clustering analysis was the apparent correlation of miRNA expression across the 14q32 locus. To test this empirically, we investigated co-expression across the locus and demonstrated a strong positive correlation between expression of 14q32 miRNAs and *MEG3*, but not with distal or proximal flanking genes/miRNAs outside of the 14q32 cluster (**Fig 3A**). This finding is consistent with *MEG3* and downstream miRNAs being expressed as a single, polycistronic transcript^61^.

**Figure 3.**
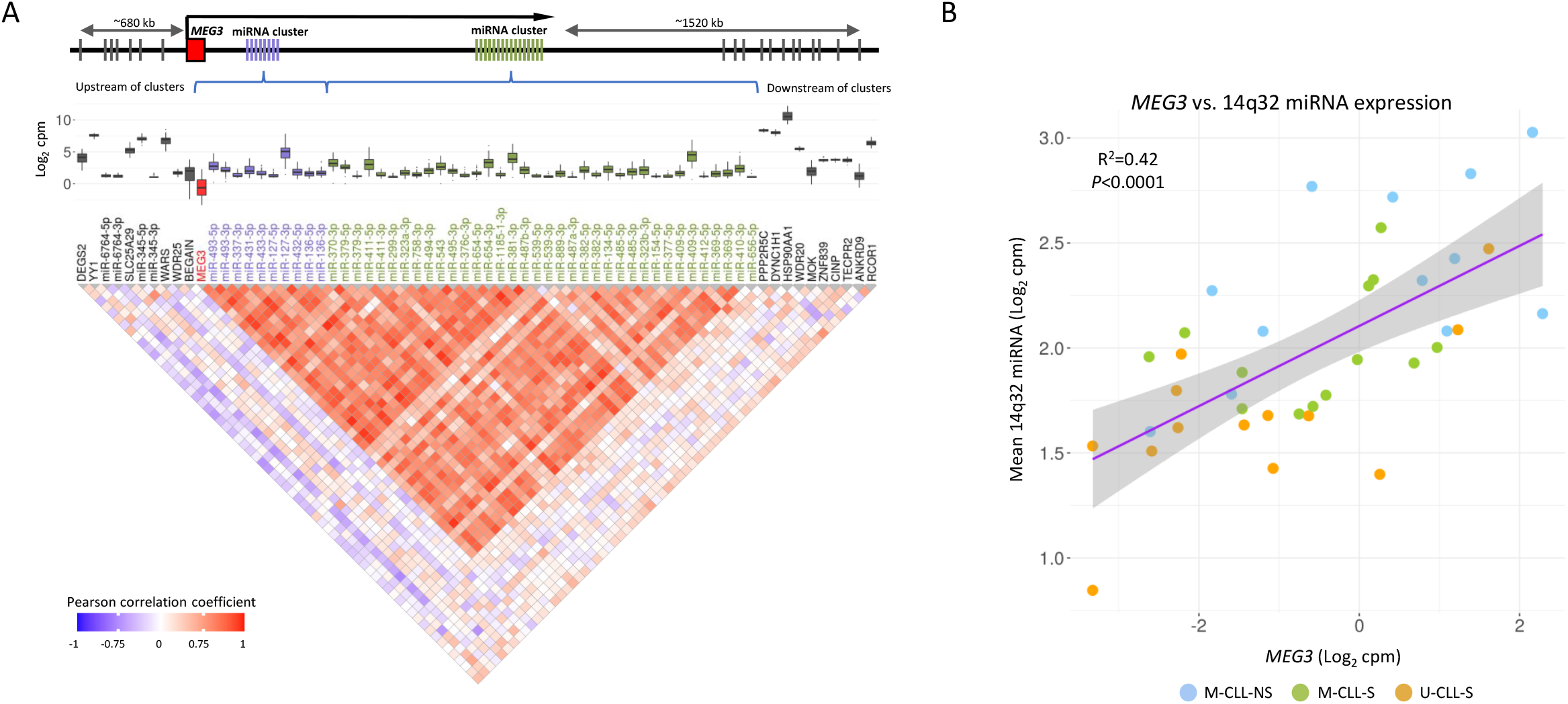
Co-expression of 14q32 locus miRNAs and mRNAs in CLL. (A) Correlation of expression of the 14q32 miRNAs and surrounding mRNAs/miRNAs depicted by a correlation plot. Red indicates a strong positive correlation between members of the 14q32 miRNA clusters and *MEG3*, blue a strong negative. miRNAs are plotted in the order that they appear in the genome, colour of miRNA name is used to separate miRNAs belonging to the 3’ and 5’ 14q32 miRNA clusters. Outside of the clusters (indicated by the genomic loci annotation) mRNA and miRNA levels do not correlate with the 14q32 miRNA cluster. Only miRNAs/mRNAs with sufficient read counts to be readily detected are depicted (see methods for inclusion criteria). (B) Correlation plot of mean 14q32 miRNA expression against *MEG3* expression.

Next, we sought to preclude the confounding impact of other clinico-biological features on the differential 14q32 miRNA expression. In contrast to multiple sclerosis, where 14q32 miRNA expression is higher in females^62^, 14q32 miRNA expression in CLL was not associated with patient sex (**Fig S6**). Whilst deletion of 14q is a rare but recurrent feature of the CLL genome, our copy number analysis detected several established recurrent copy number abnormalities (CNAs), but no deletions including 14q32^63,64^ (**Fig S7**). Although the tumour purity of all cases was high (all samples ≥77% (**Fig S1C**)), we performed confirmatory miRNA sequencing on purified CLL B-cells. We demonstrated a strong concordance between both the purified and non-purified datasets, suggesting that the 14q32 miRNA expression pattern was not driven by contaminating cells (**Fig S8A**). Specifically, the 14q32 miRNAs followed the same pattern of being downregulated in U-CLL compared to M-CLL (**Fig S8B**) and downregulated in M-CLL-S compared to M-CLL-NS (**Fig S8C**) and we observed considerable overlap between the pre-/post-purification datasets.

The regulation of imprinted gene expression at the *MEG3* locus is governed by the methylation states of two differentially methylated regions (DMRs) upstream of, and overlapping of the *MEG3* promoter (the *MEG3*-DMR and the IG-DMR respectively)^65^. Aberrant hypermethylation of the *MEG3*-DMR has been reported in various cancers^66^. However, in our data we did not observe significant differential methylation of the *MEG3* gene body, *MEG3* promoter, the *MEG3*-DMR or the IG-DMR amongst our CLL subgroups, and we did not identify substantial variation in methylation levels in this locus amongst the cohort (**Fig S9**).

### Expression of 14q32 miRNAs correlates with IGHV-dependant gene expression

Next, we aimed to identify biologically relevant network interactions within our matched miRNA and mRNA data, initially by analysing all differentially expressed mRNAs (n=498) and miRNAs (n=38). Pearson’s correlation was used to identify negatively correlated pairs of miRNA:mRNA which were intersected with databases of experimental and computationally predicted miRNA:mRNA interactions. The 198 significant, negatively correlated miRNA:mRNA pairs that were present in at least one database were taken forward as *in silico* miRNA:mRNA interactions and used to create a miRNA:mRNA interaction network for the IGHV mutational status transcriptional signatures (**Fig S10A)**. The miRNA:mRNA interaction network revealed multiple miRNAs as broad potential regulators of the CLL transcriptome, including miR-146b-5p, miR-543, miR-338-3p, miR-495-3p and miR-4476 targeting 38, 37, 15, 12 and 12 targets respectively (**Fig S10B**). Several mRNAs that were differentially expressed between U-CLL and M-CLL such as *TRIM2, TGFBR3* and *REPS2* were potentially regulated by four miRNAs each (**Fig S10C**). Characterisation of the network mRNAs by KEGG pathway membership showed mRNAs involved in Wnt (e.g., *PRICKLE2, WNT2B, WNT5B*), MAPK (e.g., *DUSP2, MAPK4*) and Ras (*GAB1*, *FGFR1, PLD1*) and BCR signalling (e.g., *GAB1, EGR2*).

Of the miRNAs identified in this network, 12 were derived from 14q32, including miR-543, miR-495-3p, miR-409-3p, miR-411-3p and miR-136-5p. The abundance of 14q32 miRNAs in the network suggested that this locus has a broad regulatory role in the CLL transcriptome. Therefore, we evaluated the potential impact of the 14q32 miRNAs on the strongest components of the IGHV-associated expression signature using the same miRNA:mRNA interaction network approach for the top 200 DEG (significant to FDR≤0.05 and Log_2_FC≥+/-1 and ranked by *P*-value) and as the 14q32 clusters appeared co-expressed/co-regulated, all 53 of the expressed 14q32 miRNAs. The analysis generated 90 interaction pairs that were used to create a 14q32 miRNA interaction network with the IGHV associated gene expression signature (**Fig 4A**). Remarkably, our miRNA:mRNA interaction network analysis showed *in silico* interaction between the 14q32 miRNAs and 49 (24.5%) of the 200 mRNAs most strongly associated with *IGHV* mutation status. We found miRNA:mRNA interaction pairs with DEG targets throughout the genome and revealed several core miRNAs with a strong potential impact on IGHV mutational status transcriptional signature through the observation of miRNAs with numerous potential target mRNAs (e.g. miR-543 was observed to have 18 mRNA interactions including *WNT5B* and *PLD1*) (**Fig 4Bi**) and some mRNAs such as *TRIM2, RNF41, GAB1* and *TGFBR3* are potentially targeted by multiple miRNAs (6, 5, 5 and 4 miRNAs respectively) (**Fig 4Bii**).

**Figure 4.**
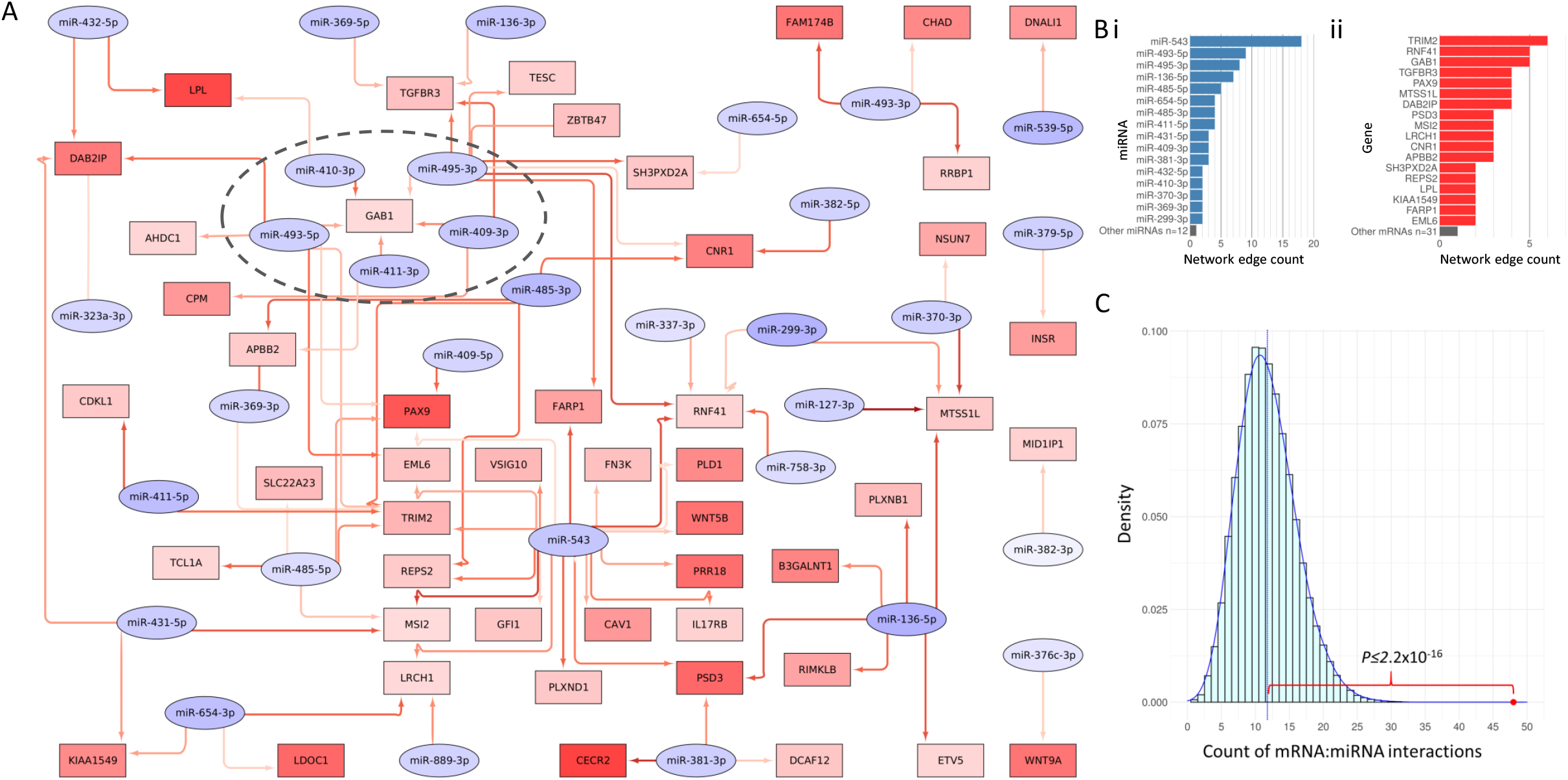
The potential impact of 14q32 miRNA on the U-CLL vs M-CLL signature. (A) A miRNA:mRNA interaction network showing the potential 14q32 miRNA regulation network of the top 200 U-CLL vs M-CLL mRNAs. Here, mRNAs are in rectangles, miRNAs are in ovals and are filled according to degree of under-expression (red) or over-expression (blue) in U-CLL vs. M-CLL. Red arrows indicate these features are negatively correlated and present in at least one putative or experimentally derived interaction database. Darker arrows indicate stronger negative correlations. *GAB1* and the network of regulatory miRNA that target it has been highlighted with a dotted oval. (B) Barplots showing the number of edges in the miRNA:mRNA interaction network for (i) each miRNA and (ii) each mRNA in the network. Features with one edge have been summarised for clarity (grey bar at bottom). (C) The number of edges in the 14q32 miRNA and top 200 U-CLL vs. M-CLL mRNA network is greater than expected by chance. We simulated 50,000 randomly selected, size matched miRNA:mRNA interactions present in miRTarget or miRDB (shown in histogram) and compared against the 48 miRTarget or miRDB edges seen in our network (red dot) using a one sample student’s t-test.

To ensure that the observation of a 14q32 network enriched for miRNA:mRNA interactions was robust, and not due to chance alone, we performed 50,000 random sampling experiments of the same number of mRNAs (n=200) and miRNAs (n=53) and compared the number of putative interactions in the TargetScan and miRDB databases for the 14q32 miRNAs network, against the 50,000 random samples (**Fig 4C**). In so doing, we demonstrated that the number of interactions associated with our 14q32 network (48 interactions) was significantly greater than those observed by chance (mean 11.75) in the randomly sampled, simulated networks (*P*≤2.2×10^−16^ from one sample Student’s t-test).

### Several 14q32 miRNAs regulate GAB1 mRNA and protein levels

A striking feature of the 14q32 network was that *GAB1*, which encodes a docking protein associated with increased BCR signalling in B cells^67^, was a putative target of 5 different miRNAs (miR-409-3p, miR-410-3p, miR-411-3p, miR-493-5p and miR-495-3p). These miRNAs were expressed in a similar manner as shown for the entire locus, i.e., high, intermediate and low mRNA expression in M-CLL-NS, M-CLL-S and U-CLL-S, respectively (**Fig 5A**). Using immunoblotting, we confirmed that GAB1 protein levels mirrored mRNA levels in our patient subgroups (**Fig 5B, Fig 5D**). As such, both *GAB1* mRNA and protein levels were inversely associated with 14q32 miRNA expression levels (**Fig 5C**).

**Figure 5.**
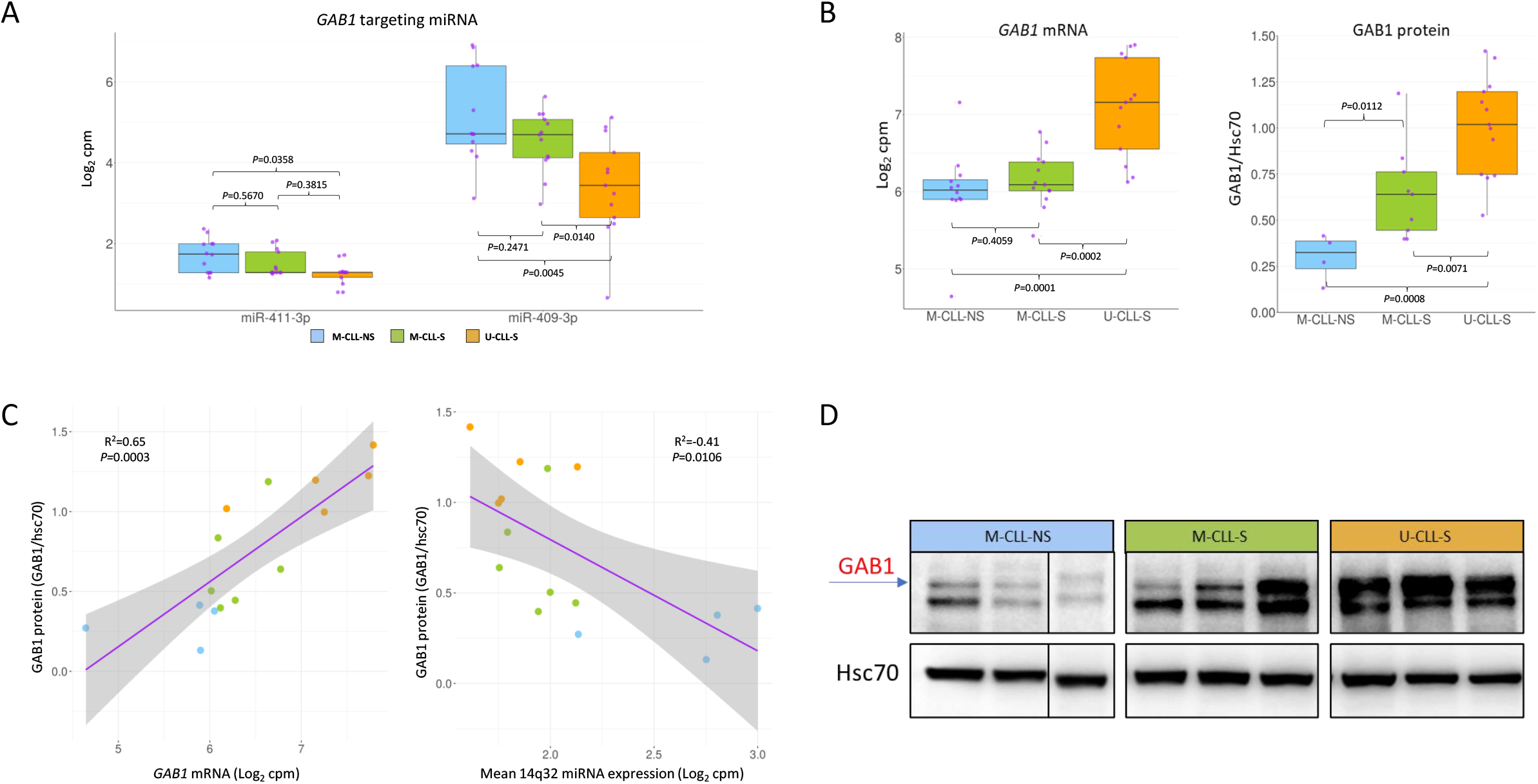
Expression of *GAB1* and *GAB1* targeting miRNA in CLL. (A) boxplots of two miRNAs predicted to interact with GAB1 *in silico* divided by CLL group. *P*-values calculated using Wilcoxon signed rank test. (B) Boxplots of *GAB1* mRNA and GAB1 protein expression for each CLL group. (C) Correlation of GAB1 protein with *GAB1* mRNA, and correlation of GAB1 protein with mean 14q32 miRNA expression. Correlation statistics generated using Pearson’s correlation coefficient. (D) Representative GAB1 immunoblots for protein level assessment.

To confirm these computationally derived, putative miRNA:mRNA interactions, we performed co-transfection experiments using pre-miR mimics, Renilla luciferase control vectors and luciferase expression vectors with a cloned *GAB1* 3’UTR in 293T cells for two miRNAs predicted to interact with *GAB1*. We chose miRNAs based on their location relative to *MEG3* and the level of support provided by the interaction prediction tools. The miRNAs miR-409-3p and miR-411-3p were able to repress activity of the *GAB1* 3’-UTR, with a 61% (*P*=0.004, Wilcoxon sum rank test) and 52% (P=0.004) reduction in luciferase reporter expression, respectively (**Fig 6**). Cross reference against target prediction algorithms showed that miR-409-3p was a previously laboratory demonstrated *GAB1* targeting miRNA^68^ whilst miR-411-3p and was predicted to interact with *GAB1* by miRTarget with a strong interaction score (85/100)^45^.

**Figure 6.**
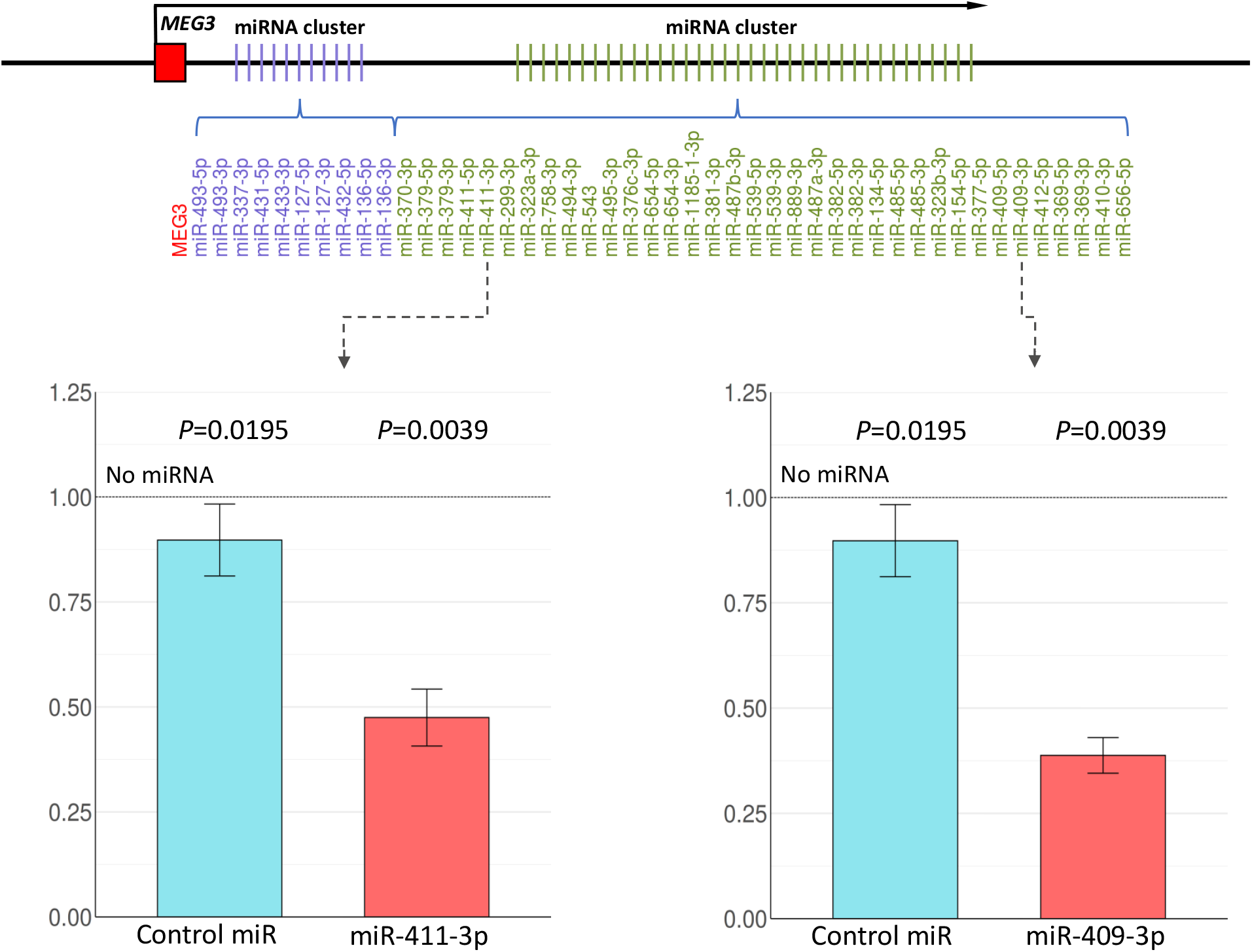
Relative luciferase intensity of luciferase expression vectors transfected alongside miRNA of interest or control miRNA. Relative luciferase expression ratios were calculated by dividing the control/test miRNA luciferase levels by a no miRNA control. Ratios were tested statistically using one sample Wilcoxon signed rank tests to mu of 1 (i.e. the no miRNA control). Error bars indicate standard deviation to mean. Upper annotation depicts the relative position of selected miRNA in the 14q32 miRNA clusters and relative to *MEG3*. Annotation is not to scale.

## Discussion

Detailed analysis of the CLL tumour cell has revealed the landscape of its genome, the relationship between transcriptome and epigenetic regulation, and the structure and function of the BCR. Despite these expansive studies, the extent to which transcriptional regulation reflects the innate pre-/post-germinal centre origin of U-CLL and M-CLL or BCR signalling competence remains unanswered. As such, we employed high throughput molecular approaches to correlate genome-wide miRNome and transcriptome signatures and DNA methylation patterns with IGHV and BCR signalling characteristics. In doing so, we identified two miRNA clusters located in 14q32 that strongly associated with IGHV status and were tightly co-expressed alongside the lncRNA *MEG3*. To ensure the validity of our findings, we precluded the impact of 14q deletion and patient sex and limited the effect of laboratory manipulation by focussing our transcriptome/miRNome analyses on non-purified CLL cells from high purity samples. To curtail the impact of contaminating germline cells, we validated our finding on CLL cells after the depletion of non-tumour cells. We demonstrated that miRNAs within these clusters were predicted to have an expansive regulatory role in IGHV-dependent transcriptional control, exemplified by the targeting of *GAB1* that was confirmed by *in vitro* functional assays.

Our U-CLL vs. M-CLL mRNA expression signature showed broad agreement with published works^22,53,69^, despite historical changes to miRNA nomenclature, technical differences and a degree of inconsistency between studies; for example, data is inconclusive on the relationship between miR-155-5p expression and IGHV mutation status^27,28,51,52,70^. In our study, miR-155-5p expression was not associated with IGHV status in either purified or non-purified CLL tumour cells, which may reflect the composition of our cohort as chromosomal aberrations linked to miR-155-5p expression such as trisomy 12 and deletion of 17p^54^ were scarce in our cohort. The inability of previous miRNA studies to identify dysregulation of the 14q32 locus is likely to be the result of the paucity of 14q32 miRNA coverage on microarray/qPCR platforms and the enrichment of the high 14q32 miRNA expression M-CLL-NS cases in our cohort.

Whilst our study is the first to report a role of the 14q32 cluster miRNAs in CLL subtypes, they have been extensively studied in solid tumours and other haematological neoplasms and are reported to have tumour suppressor roles in melanoma (miR-376a/c), papillary thyroid (miR-654-3p), colorectal (miR-409-3p) and renal cell cancer (miR-411-5p)^68,71–73^. In contrast, high-expression of 14q32 miRNAs have been associated with inferior outcome in lung cancer and acute myeloid leukaemia, suggesting that the cluster may have cell-type specific functions^74,75^. In splenic marginal zone lymphoma (SMZL) several 14q32 miRNAs are down-regulated compared to normal B cells and CLL^76^, suggesting the cluster may have a more expansive role in B-cell tumours. The strong correlation observed amongst individual 14q32 miRNAs and the *MEG3* lncRNA supports mouse studies reporting expression from a ^~^200 kbp polycistronic transcript originating from the *MEG3* TSS^61,77^. *MEG3* was the first tumour suppressor lncRNA identified, has been demonstrated to have a pathogenic role in a number of cancer models^78,79^, and is down-regulated in a number of primary tumours in comparison to matched normal tissues^80,81^. Further analysis of *MEG3* was hindered by the low expression level and it was difficult to delineate *MEG3* function from miRNA function in CLL given the strong co-expression observed.

*MEG3* and the 14q32 miRNAs are located within the *DLK1-DIO3* genomic imprinted region, controlled by the paternal/maternal allele specific methylation of the two differentially methylated regions (DMR), the IG-DMR 13 kbp upstream of the promoter and the *MEG3*-DMR overlapping the *MEG3* promoter^65^. Aberrant hypermethylation of the *MEG3*-DMR is a feature of lymphoid and myeloid neoplasms^66,82^, for example in acute promyelocytic leukaemia where loss of IG-DMR imprinting is associated with increased *MEG3* and 14q32 miRNA expression^83^. In our cohort, we did not identify differential DMR methylation. However, the arrays used had poor coverage of the IG-DMR, limiting our analysis and emphasising the need for analysis of this cluster at greater resolution to draw definitive conclusions.

In addition to differential expression of the entire 14q32 locus between M-CLL and U-CLL, we observed a stepwise increase from U-CLL-S to M-CLL-S and M-CLL-NS, that was apparent in both purified and non-purified samples. This observation was supported by our unsupervised clustering analysis that showed three levels of 14q32 miRNA expression levels, broadly associated with our three IGHV/BCR signalling groups. High BCR signalling capacity has been demonstrated to be a poor prognostic indictor, independent of IGHV mutation status^12^. As such, the potentially inverse association between 14q32 miRNA expression and prognosis in our CLL subgroups is notable in that it would coincide with the tumour suppressor potential of the 14q32 miRNAs and *MEG3* lncRNA. Our study lacked the power to discriminate survival differences associated with 14q32 miRNA expression independently of other factors, but it would represent an area of exciting potential for future work.

When we classified our patients by DNA methylation epitype we showed the expected enrichment of m-CLL and n-CLL in the M-CLL and U-CLL subgroups, respectively. Whilst there was some evidence of an enrichment of m-CLL cases in the M-CLL-NS group, i-CLL cases in the M-CLL-S group and n-CLL cases in the U-CLL-S group, there was not a strong association between epitype and 14q32 miRNA expression. The analysis of a larger CLL cohort would be needed to establish the exact relationship between epitype, IGHV status, BCR signalling competence and 14q32 miRNA expression, but our preliminary analysis suggests that 14q32 expression is more associated with IGHV and BCR signalling than epitype.

Remarkably, our miRNA:mRNA interaction network analysis showed that the 14q32 miRNAs have a potential for broad perturbation of entire networks of miRNA-mRNA interactions, where 49 (24.5%) of the 200 strongest features of the IGHV associated transcriptional signature may be regulated by 14q32 miRNAs. The limitations of our approach include an assumption of interaction for negatively correlated mRNAs and miRNAs where the interaction is plausible from miRNA:mRNA interaction databases, and our approach is clearly only able to identify the consequences of mRNA destabilisation, as any impact on translation would not be apparent at the miRNome and transcriptome level. We were able to confirm the mechanistic link between several 14q32 miRNAs and regulation of the *GAB1* gene which encodes a docking protein that when phosphorylated can increase BCR signalling in B cells ^67^. Therefore, in addition to the previously reported ability of miR-150-5p to reduce *GAB1* expression in CLL cells^28^, we now show that two further miRNAs, miR-411-3p and miR-409-3p, located at 14q32 also regulate *GAB1* and associate with IGHV and BCR signalling status.

Questions remain about the ability of 14q32 miRNAs to regulate the expression of *GAB1 in vivo* and their role in other B-cell malignancies, during B-cell development and in response to treatment, all of which represent promising opportunities for future work. We speculate that the dysregulation of multiple miRNAs could provide precise regulatory control, reduce the degree of redundancy in the epigenetic regulation of key mRNAs, or have an additive impact on transcriptional regulation but these hypotheses remain to be confirmed. Our data hint at an association between 14q32 miRNA expression level and survival that if officiated could reveal 14q32 miRNA quantification as prognostic biomarkers. Further investigation and validation of the methylation dependent regulation of the 14q32 locus in CLL, perhaps using long read technologies that offer high resolution DNA sequencing, methylation status and phasing of methylated CpGs across the locus would highlight the potential for demethylating agents such as decitabine to relieve repression of 14q32 miRNAs and *MEG3* in high risk CLL subsets.

Our data illustrates the critical role of miRNA-mediated regulation in the pathobiology of CLL, showing that the 14q32 miRNAs have a putative regulatory role in IGHV-associated transcription, with functional evidence of a mechanistic interaction with *GAB1*. Given that these naturally produced molecules and their levels can be readily regulated with miRNA mimics or miRNA/antagomiRs, their therapeutic manipulation may have implications for more effective deployment of existing targeted agents.

## Acknowledgements

We are grateful to Dr Kathy Potter at the Faculty of Medicine Tissue Bank (Cancer Sciences, University of Southampton) for the processing and storage of the primary CLL specimens. This work was funded by Cancer Research UK (ECRIN-M3 accelerator award C42023/A29370, Southampton Experimental Cancer Medicine Centre grant C24563/A15581, Cancer Research UK Southampton Centre grant C34999/A18087, and programme C2750/A23669). The authors acknowledge the use of the IRIDIS High Performance Computing Facility, and associated support services at the University of Southampton, in the completion of this work.

## Individual contributions

GP, JS, AJS, DB & LS designed the study, DB, JS, AJS & GP wrote the manuscript, DB, LS, AJS & JW performed data analysis, DB, LS, KRB, LK, AR, MDB & PH performed laboratory work. All authors reviewed the manuscript.

