## Supplementary Figures for "Network analysis reveals a major role for 14q32 cluster miRNAs in determining transcriptional differences between IGHV-mutated and unmutated CLL"

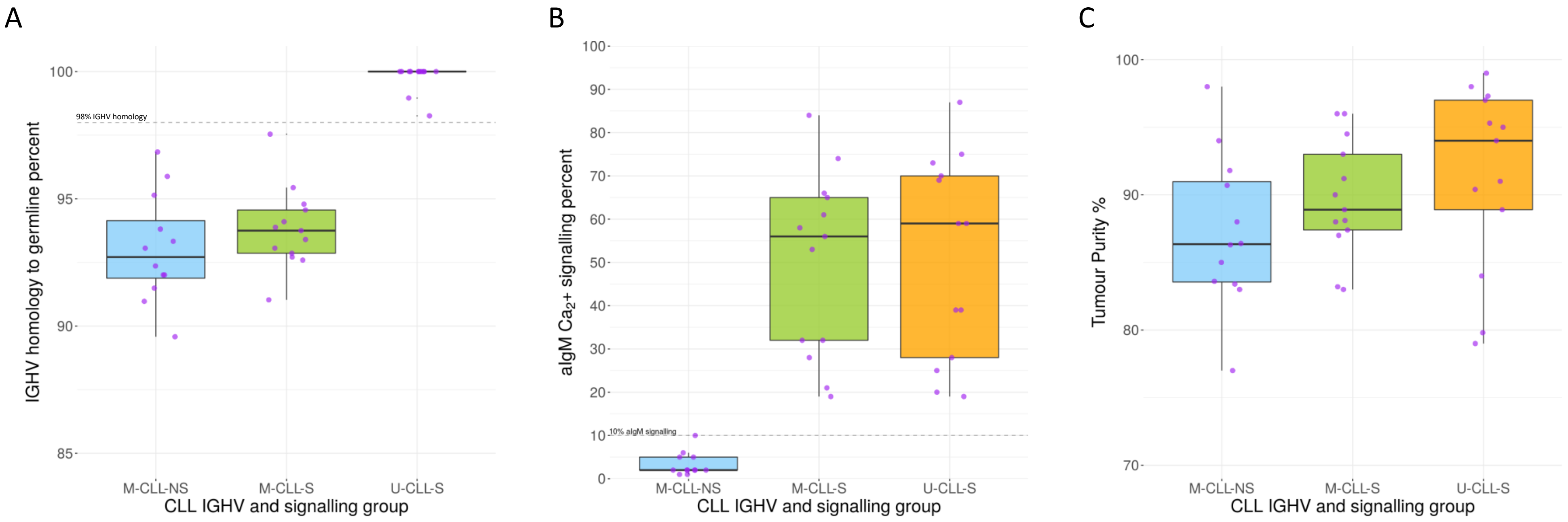


**Supplementary Figure 1. Phenotypic characterisation of the CLL cohort.** (A) A boxplot showing the IGHV percent homology to germline of the three CLL groups. (B) Boxplot showing the percentage of cells exhibiting $Ca^{2+}$ flux as a signalling response to anti-IgM ligation for the three CLL groups. (C) Boxplot showing the percentage of tumour cells (i.e., tumour purity) in each sample for the three CLL groups. In all plots, purple points represent individual samples.

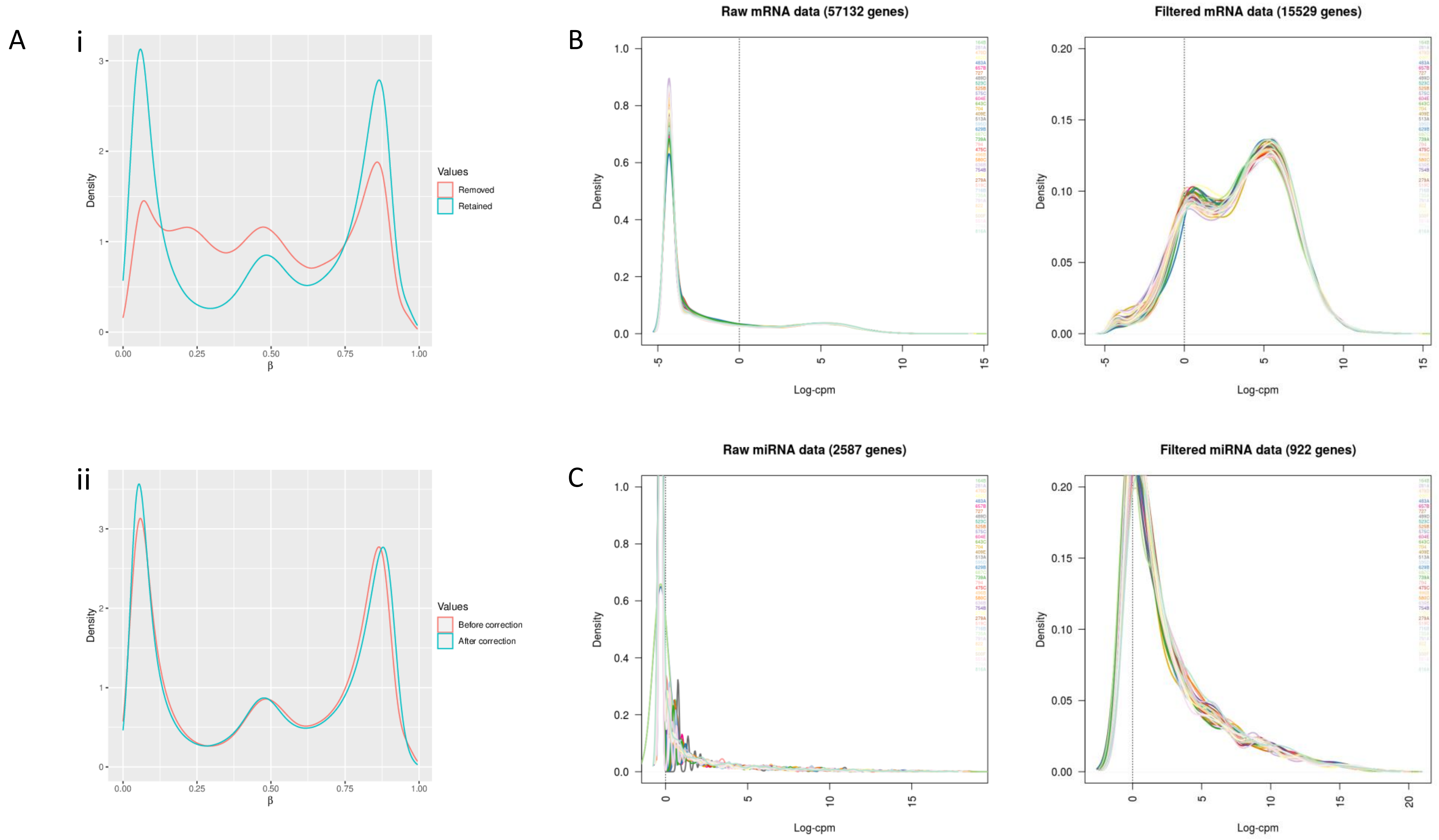


**Supplementary Figure 2. Filtering and normalisation of DNA methylation array data mRNA/miRNA sequencing data.** (A) i) Beta Density plot of retained/removed probes, (ii) beta density plot before and after SWAN normalisation. (B,C) cpm density plot before and after filtering of the miRNA and mRNA data. Filtering was performed in order to minimise the number of features with low read counts.

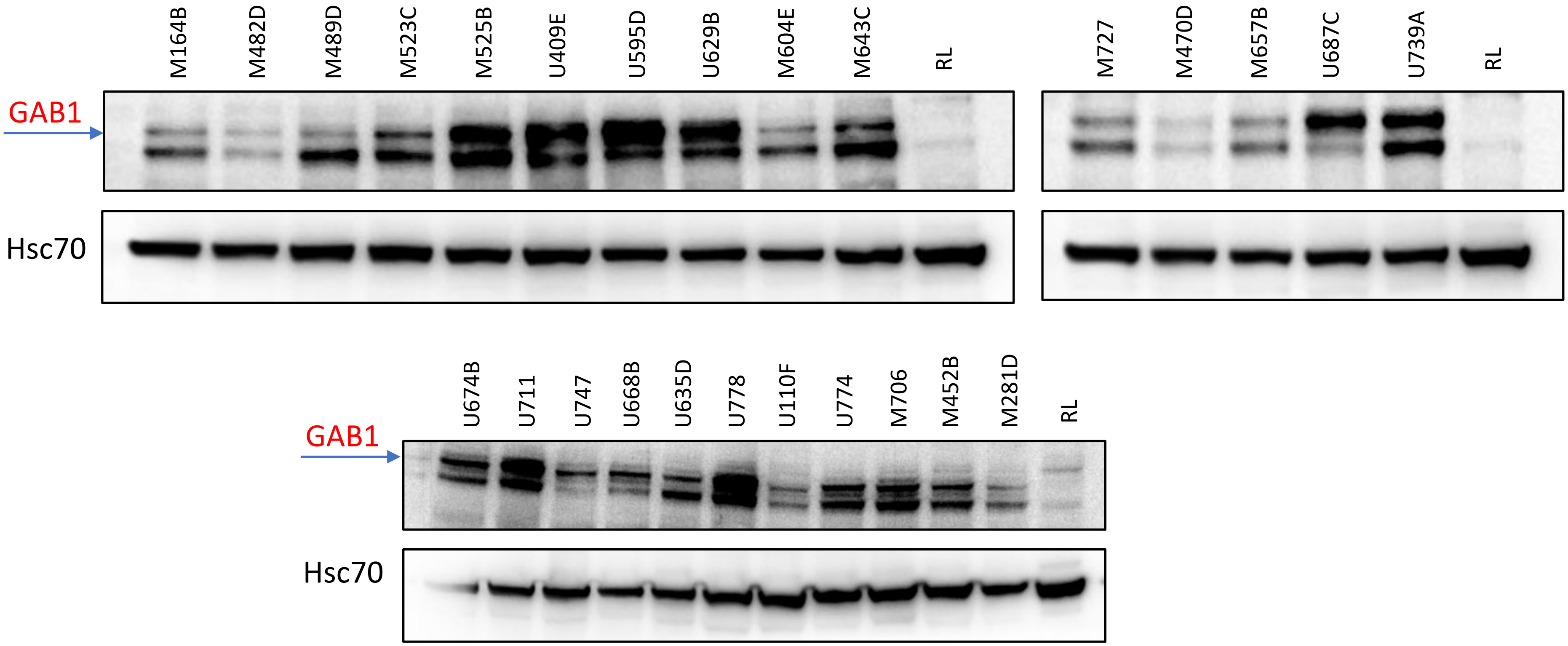


**Supplementary Figure 3. Representative GAB1 immunoblots for protein level assessment.** The top bands are GAB1, the second bands are GAB2/GAB3, also shown is the Hsc70 control.

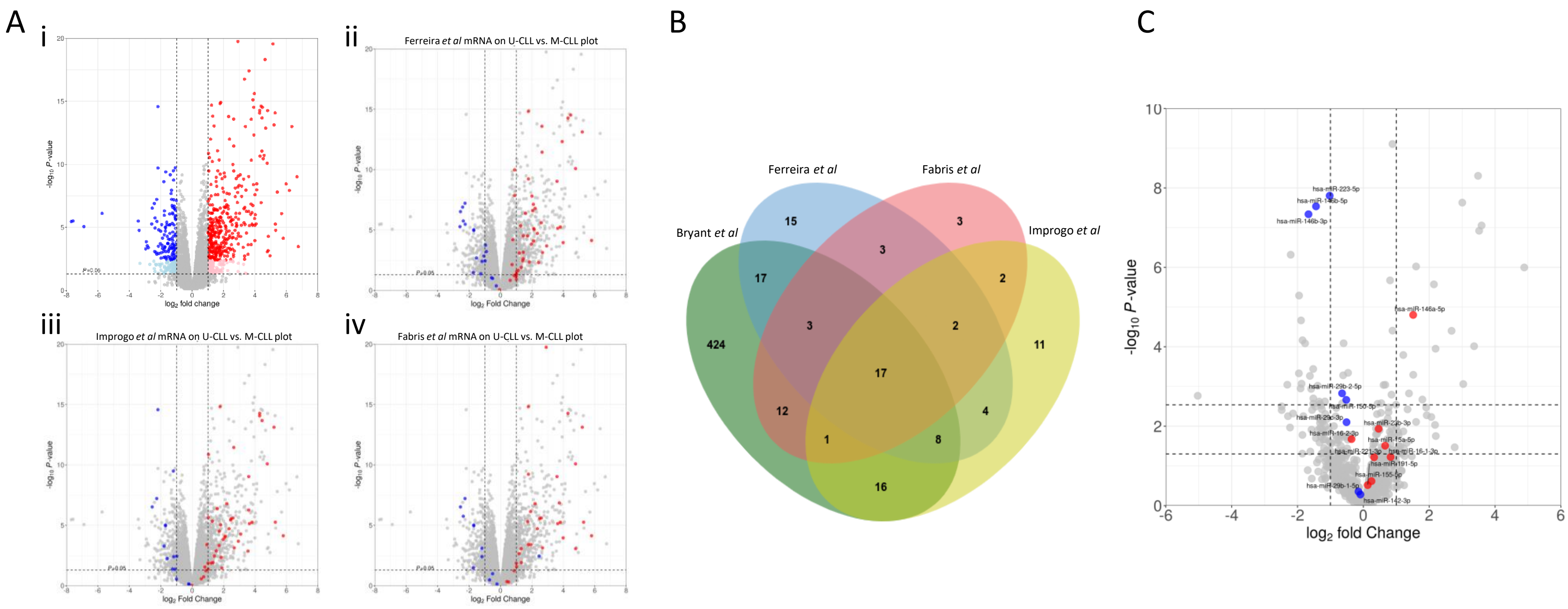


**Supplementary Figure 4. Analysis of the overlap between our U-CLL vs. M-CLL differentially expressed gene lists against public data.** (A) The U-CLL vs. M-CLL volcano plot from this study colour coded by the differential expression results from 3 published datasets. (A) i) The U-CLL vs M-CLL differential gene expression results from this study. ii) The differential expression results from this study, highlighted according to the differential expression result in Ferreira *et al* 2014. iii) The differential expression results from this study, highlighted according to the differential expression result in GEO dataset GSE69034 (Improgo *et al* 2019). iv) The differential expression results from this study, highlighted according to the differential expression result in GEO dataset GSE38611 (Fabris *et al* 2013). (B) A venn diagram showing the number of differentially expressed genes in common between this study and the three published datasets. (C) Overlap of a number of key miRNAs described as differentially expressed in U-CLL compared to M-CLL (Mraz *et al* 2009, Negrini *et al* 2014).

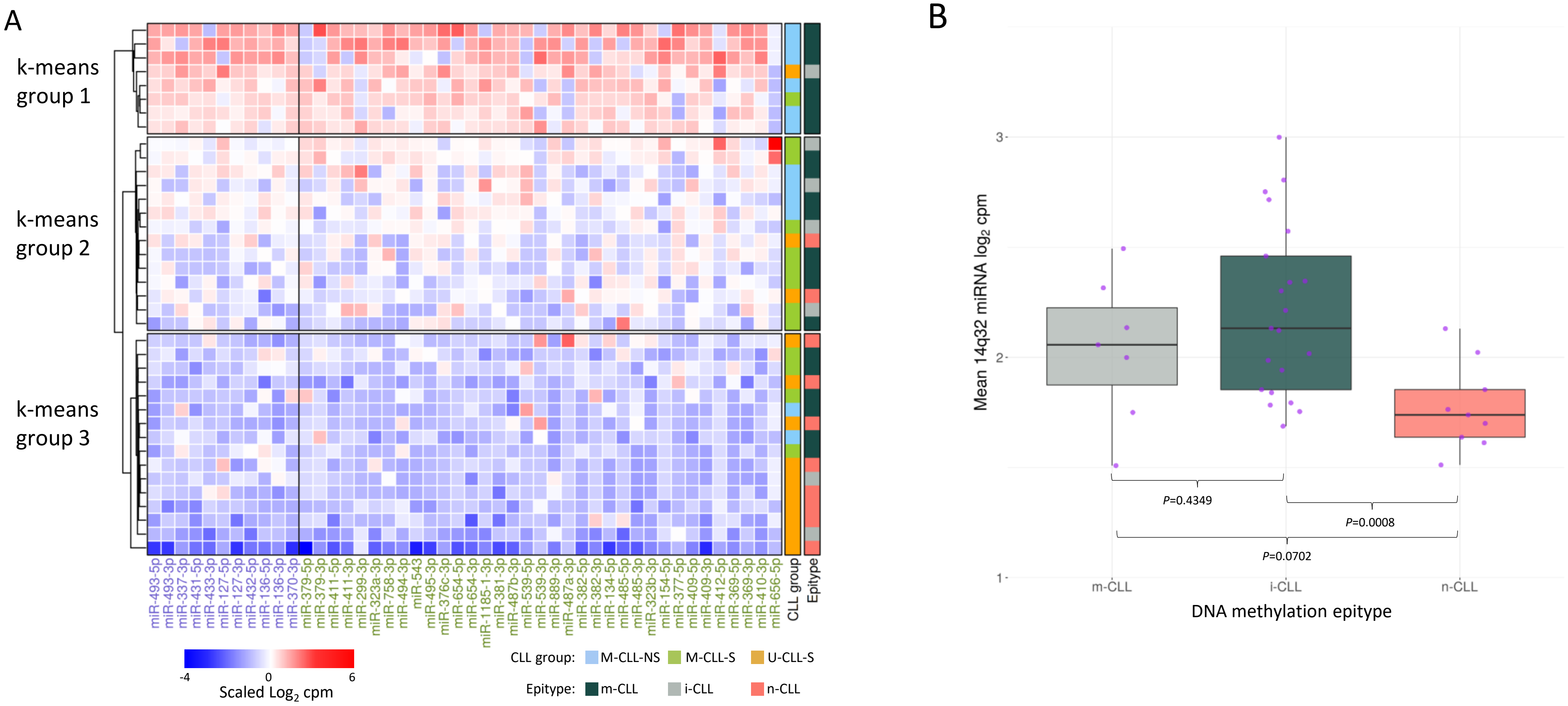


**Supplementary Figure 5. Heatmaps and clustering analysis of 14q32 miRNA expression and association with CLL subgroup and epitype.** (A) Heatmap showing expression of 14q32 miRNA. High expression is indicated by red boxes, low expression is indicated by blue boxes. The x-axis shows miRNA names in the order they appear in the genome, colour is used to separate miRNA belonging to the 3' and 5' 14q32 miRNA clusters. Only miRNAs with sufficient read counts to be included in statistical analysis are depicted (see methods for inclusion criteria). The y-axis is split into three groups as determined using unsupervised k-means clustering (k-means groups 1-3). Hierarchical clustering shown in dendrograms of both plots. Row annotation shows CLL subgroup and epitype. (B) Boxplot of mean 14q32 miRNA expression by DNA methylation epitype (Kulis *et al* 2012). Purple points are samples assigned to each epitype.

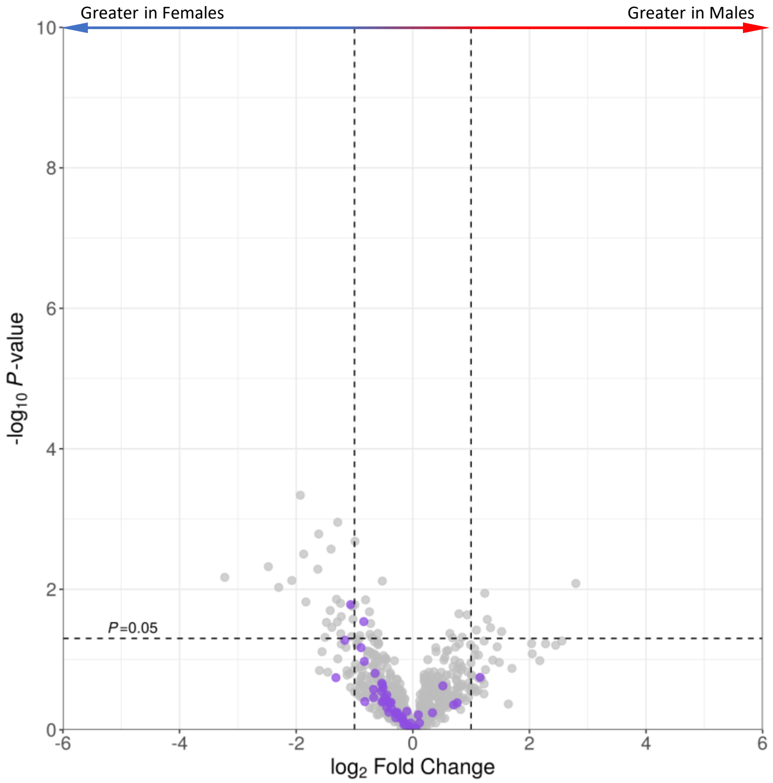


**Supplementary Figure 6. The effect of patient sex on 14q32 miRNA expression.** The male vs. female volcano plot colour coded by 14q32 miRNAs (purple points) and non 14q32 miRNAs (grey points)

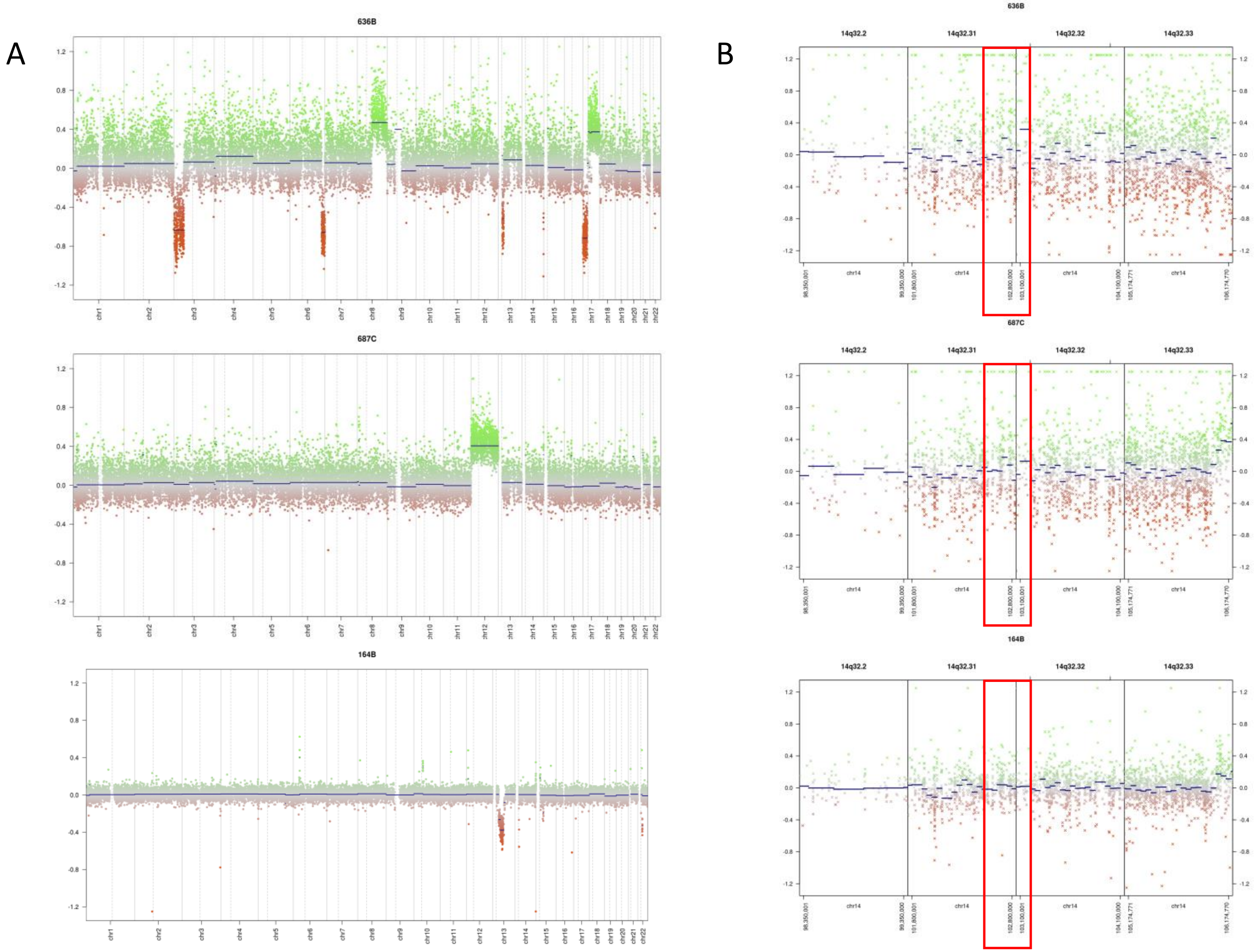


**Supplementary Figure 7. Copy number profiles derived from EPIC methylation array intensities for three representative samples from the CLL cohort.** Here, the Y-axis indicates copy number changes compared to a cohort of normal B cells, with values greater than 0 indicating a copy number gain and values below 0 indicating a copy number loss. Coloured points represent bins of CpGs whereas the blue lines indicate called and segmented CNV profiles. (A) Genome wide copy number profiles for three representative samples of the CLL cohort showing various CNVs but no evidence of 14q32 deletion. (B) Higher resolution view of the 14q32 region in three representative CLL samples, the 14q32 miRNA cluster locus is indicated by the red boxes. Noise towards the right is indicative of IGH gene selection in CLL and controls.

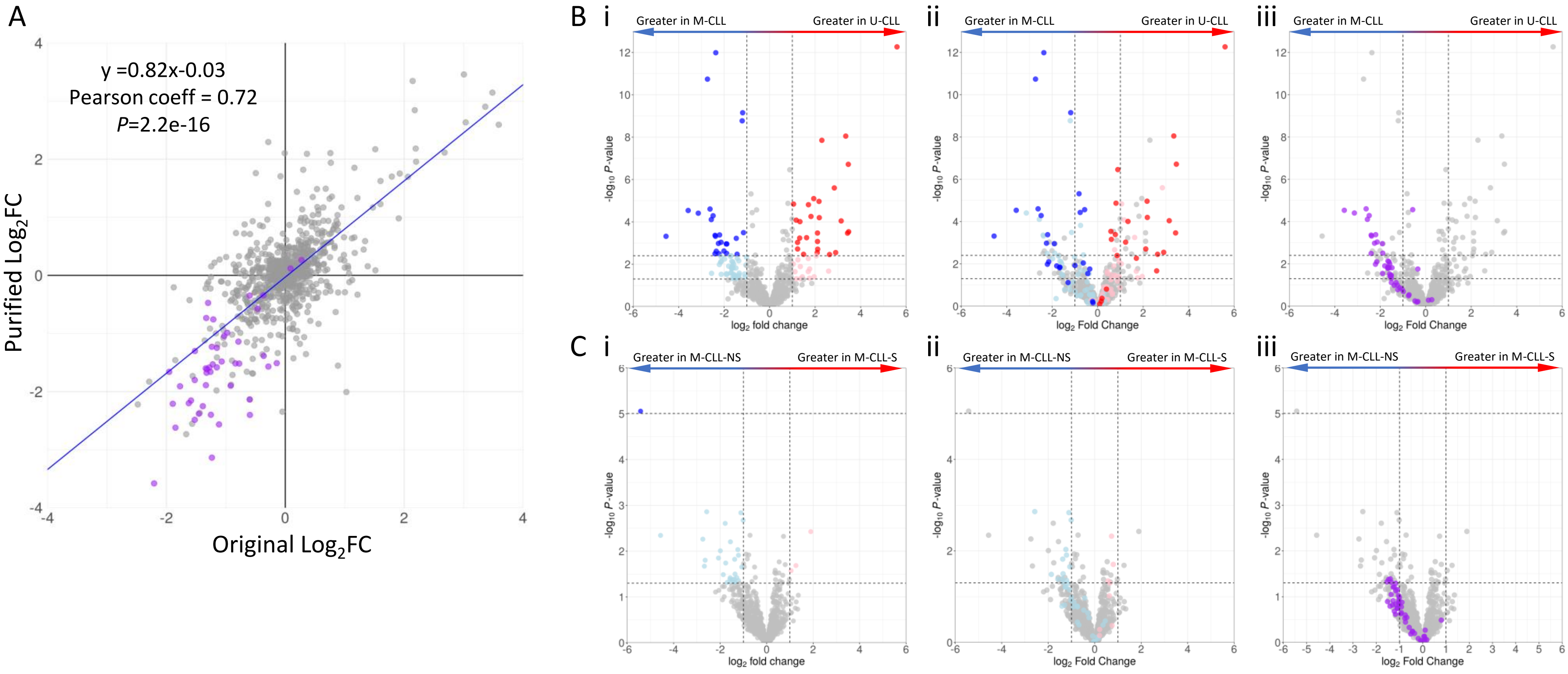


**Supplementary Figure 8. Comparison of differential miRNA expression in non-purified and purified CLL cells.** (A) Scatterplot comparing U-CLL vs. M-CLL $Log_2FC$ in non-purified CLL cells (x-axis) and purified CLL cells (y-axis). Purple points are 14q32 miRNAs. (B) U-CLL vs. M-CLL differential miRNA expression comparisons in purified and non-purified CLL cells using volcano plots. i) Differential miRNA expression in purified CLL cells. ii) How the DEMs in non-purified CLL cells behaved in the purified CLL cells. iii) The 14q32 miRNAs (purple points) were downregulated in purified CLL cells. (C) M-CLL-S vs. M-CLL-NS differential miRNA expression comparisons in purified and non-purified CLL cells using volcano plots. (i) Differential miRNA expression in purified CLL cells. (ii) How the miRNAs in non-purified CLL cells behaved in the purified CLL cells. iii) The 14q32 miRNAs (purple points) were downregulated in purified CLL cells. Each point represents a miRNA and the fold change and P-value for differential expression. Grey points were not differentially expressed, points to the right of zero are over expressed, points to the left are under-expressed. Light blue/red are differentially expressed at *P*=0.05 whilst dark blue/red are differentially expressed at FDR=0.05 after correcting for multiple testing.

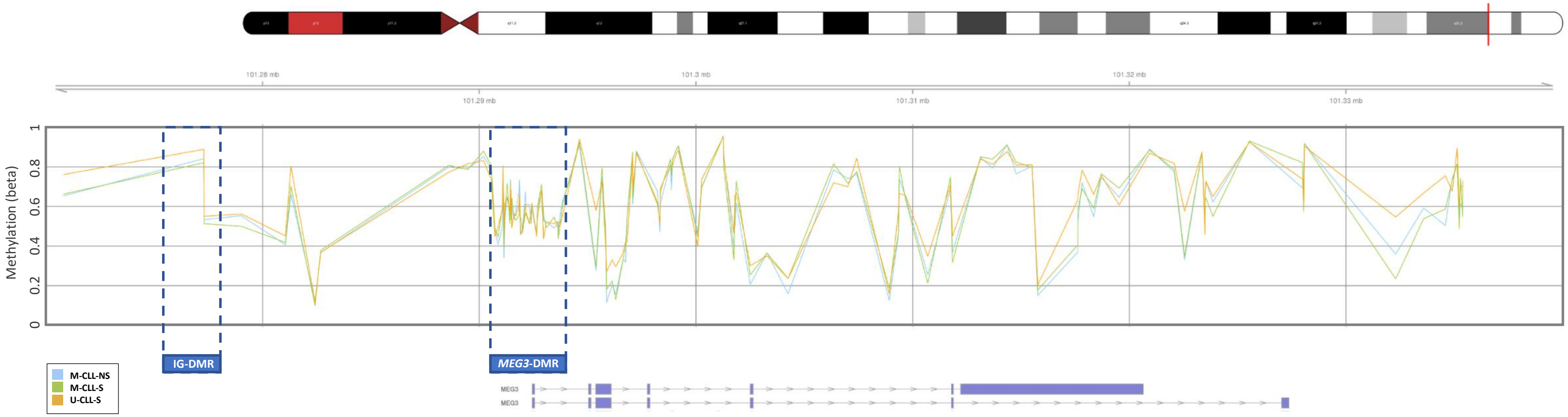


**Supplementary Figure 9. DNA methylation of the *MEG3* locus in CLL.** Mean DNA methylation as determined by EPIC methylation array analysis is shown on an individual CpG basis for each of the three CLL groups (i.e. M-CLL-NS, M-CLL,S and U-CLL-S). The IG-DMR and *MEG3* DMR represent regions of differential methylation between maternal and paternal alleles during developmental genomic imprinting. There is no evidence of a CLL group specific loss of imprinting in either DMR with Beta values in both regions averaging approximately 50%. Connection lines between CpG methylation values are drawn to aid interpretation and do not infer or describe a relationship beyond adjacency.

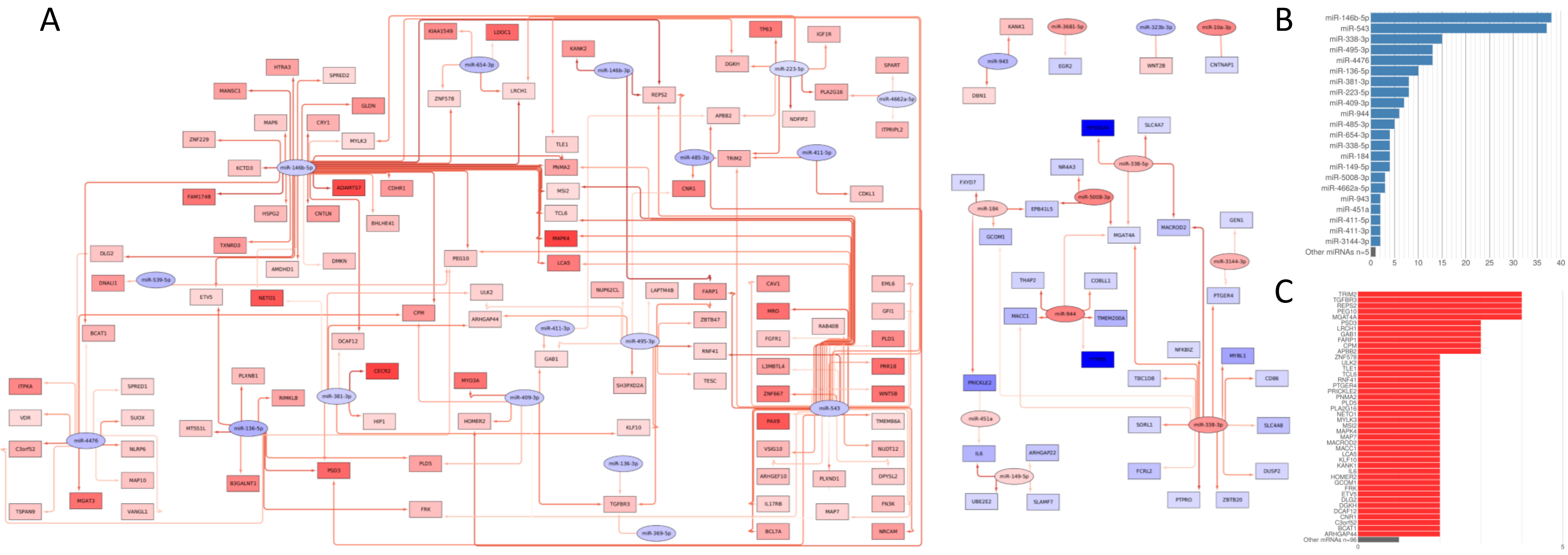

**Supplementary Figure 10. The potential impact of differentially expressed U-CLL vs. M-CLL miRNAs on the differentially expressed U-CLL vs. M-CLL mRNA signature.** A miRNA:mRNA interaction network showing the potential miRNA regulation network of differentially expressed U-CLL vs. M-CLL mRNAs. All miRNAs and mRNAs are differentially expressed at $Log_2FC\geq+/-1$ and FDR≤0.05. Here, mRNAs are in rectangles, miRNAs are in ovals and are filled according to degree of over-expression (red) or under-expression (blue) in U-CLL vs M-CLL. Red arrows indicate these features are negatively correlated and present in at least one putative or experimentally derived interaction database. Darker arrows indicate stronger negative correlations.
