## Supplementary Methods for "Network analysis reveals a major role for 14q32 cluster miRNAs in determining transcriptional differences between IGHV-mutated and unmutated CLL"

### Patient cohorts and sample characteristics

An overview of the cohort and methods applied in this study are depicted in Figure 1A (**Fig 1A**). Thirty-eight basal tumour samples, obtained from CLL patients diagnosed using the iwCLL guidelines[1] and managed at Southampton General Hospital (Southampton, UK), were selected based on IGHV mutational status using established cut-offs[2] (**Fig S1A**), and BCR signalling capacity with >10%/≤10% anti-IgM-induced $Ca^{2+}$ mobilisation thresholds (**Fig S1B**) determined as described previously[3]. Informed consent was obtained in accordance with the declaration of Helsinki and the study was approved by our regional research ethics committee. We focused on high purity (mean 89.5%, range 77%-99%) (**Fig S1C**), but non-purified CLL cells for the initial transcriptome and miRNA sequencing and used purified CLL cells (Miltenyi B-CLL Isolation kit) for DNA methylation analysis and confirmatory miRNA sequencing. CLL cases were divided into three subgroups of BCR-signalling competent U-CLL (U-CLL-S, n=13) and M-CLL (M-CLL-S, n=13) and BCR-signalling deficient M-CLL (M-CLL-NS, n=12). Due to limitations in sample material, GAB1 protein immunoblotting was performed on a subset of our main cohort comprising U-CLL-S (n=5), M-CLL-S (n=6) and M-CLL-NS (n=4), and additional CLL samples with BCR signalling and IGHV data available, including U-CLL-S (n=8) and M-CLL-S (n=3).

### mRNA and miRNA sequencing

mRNA sequencing was performed on RNA extracted from $1x10^6$ CLL cells purified with the Qiagen RNeasy mini prep kit (Qiagen, Hilden, Germany) and libraries were prepared using TruSeq RNA kits (Illumina, Hayward, CA, USA). Sequencing was performed in The Garvan Institute of Medical Research (University of New South Wales, Sydney, Australia) and by the Oxford Genomics Centre (Oxford, UK) using an Illumina HiSeq2500 and HiSeq4000 (Illumina, Hayward, CA, USA), respectively, to a mean of $5.6x10^7$ (range 4.4 $x10^7$-8.0 $x10^7$) 75 bp SE and $8.9x10^7$ (range 6.7 $x10^7$-$1.2x10^8$) 75 bp PE reads, respectively. Our sample subgroups were balanced across sequencing runs to improve batch correction. Raw mRNA sequencing data in the form of fastq, trimmed of sequencing adaptors using FastX toolkit (http://hannonlab.cshl.edu/fastx_toolkit/), was quality checked using FastQC (Babraham Bioinformatics, Cambridge, UK) then aligned to the hg38 reference genome using HISAT2[4] (a mean of 96.0% of reads aligned (range 92.4%-98.8%). Read counts per gene were calculated using HTseq-count[5] against the Ensembl GRCh38 v94 GTF[6].

miRNA library preparation was performed on total RNA extracted with miRNeasy kits (Qiagen, Hilden, Germany) using Illumina TruSeq small RNA library kits (Illumina, Hayward, CA, USA). Sequencing of libraries was performed by two centres, 18 samples were sequenced by Q2 Solutions (Q2 solutions, Valencia, CA, USA) to a mean of $8.3x10^6$ (range $7.0x10^6$-$9.9x10^6$) SE 50 bp reads per sample using an Illumina HiSeq 4000 (Illumina, Hayward, CA, USA) and 21 samples were sequenced by Oxford Genomics Centre (Oxford, UK) to a mean of $1.5x10^7$ (range $8.3x10^6$-$1.9x10^7$) 50 bp SE reads an Illumina HiSeq 2500 (Illumina, Hayward, CA, USA). Our sample subgroups were balanced across sequencing runs to improve batch correction. Subsequent, confirmatory miRNA sequencing was performed to preclude any potential contribution of contaminating T-cell/monocytes to our findings, processed as before, but from CLL purified using the Miltenyi B-CLL Isolation kit according to kit instructions with LS Columns (Miltenyi, Bergisch Gladbach, Germany). Subsequent sequencing was performed by Oxford Genomics Centre (Oxford, UK) by sequencing to 7-$10x10^6$ SE 50 bp reads per sample using Illumina TruSeq small RNA library preparation kits and an Illumina HiSeq 4000 (Illumina, Hayward, CA, USA). Confirmatory sequencing was performed as a single batch. miRNA data in the form of fastq were quality checked using FastQC, then aligned to the hg38 reference genome using the BWA -aln algorithm of BWA v0.7.12[7]. miRNA read counts were calculated using HTseq-Count against miRbase v21[8].

### EPIC DNA methylation array

DNA methylation was assessed in purified CLL tumour cells (Miltenyi B-CLL Isolation kit), using EPIC DNA methylation arrays (Illumina, Hayward, CA, USA) performed by the Statens Serum Institute, Denmark. Data was analysed using RnBeads v2.93[9] from raw intensity data through to import, annotation against hg19, quality control, SWAN normalisation, differential methylation analysis and output beta/M values. Following removal of SNP enriched probes (17,371) and unreliable probes using the GreedyCut method (17,302), 832,222 probes were used for the analysis (**Fig S2A**). Differential methylation at site and regional level was calculated using the limma method with a minimum delta beta of 0.25 and adjusted *P*-Value of 0.05 (Benjamini Hochberg method, FDR). Conumee[10] was used to produce copy number profiles from mean intensity signals. In house EPIC array data from a cohort of copy number neutral, normal B cell subsets from 3 individuals were used as a baseline control. DNA methylation epitypes were determined using the process described by Kulis et al[11], using the 1,502 of the 1,649 CpGs described in the epitype methylation signature that are present in the Illumina EPIC array. Consensus clustering was performed with 80% subsampling of both items and features, max k=10 and bootstrapped 10,000 times[12]. All other analysis were performed in R v.3.6.1 (R Development Core Team 3.0.1., 2013) using custom code.

### Immunoblotting

SDS-PAGE was performed on $3x10^6$ lysed CLL cells and run with equal protein loading on 10% Nu-PAGE Bis-Tris gel (Invitrogen, Waltham, MA, USA) with MOPS buffer (Invitrogen, Waltham, MA, USA). Protein quantitation was evaluated using the Bio-Rad Protein Assay (Bio-Rad Laboratories Inc, Hercules, CA, USA.) and the blots stained with the following primary antibodies; rabbit anti-GAB (Cell Signaling Technologies, Danvers, MA, USA) and mouse anti-Hsc70 (Santa Cruz Biotechnology Inc, Dallas, Texas, USA). Secondary antibodies were horseradish peroxidase-conjugated anti-rabbit/anti-mouse (Dako, Santa Clara, CA, USA). Images were captured using the ChemiDoc-It Imaging System and quantified using ImageJ (http://imagej.nih.gov/ij/). The GAB antibody recognised other isoforms of GAB and GAB1 was represented by the upper band on the immunoblot as determined by size (110 KDa). GAB1 expression was normalized to the expression of Hsc70 for each sample.

### miRNA transfections

miRNA activity was analysed by co-transfecting 293FT cells (ThermoFisher, Leicestershire, UK) with Lipofectamine 2000 (ThermoFisher, Leicestershire, UK) and a human *GAB1* 3'-UTR reporter plasmid (containing 3770 bp immediately downstream of the end of the GAB1 ORF cloned into pMirTarget, Origene), a control Renilla luciferase plasmid (Promega) and pre-miR miRNA mimics or pre-miR control 1 (all ThermoFisher, Leicestershire, UK) (Fig 3A & Fig 3B). Cells were also transfected with the plasmids but without the pre-miR as an additional no miRNA control. After 24 hours, luciferase activity was quantified using the dual-glo luciferase assay system (Promega, Southampton, UK).

Firefly luciferase was normalised using Renilla luciferase values from the same well and normalised values for control transfected cells (no pre-miR) were set to 1.0.

Data analysis

Filters were applied to both mRNA and miRNA data to remove low expression genes and miRNA (only features with ≥1 cpm in ≥3 (mRNA) or ≥2 (miRNA) samples were retained) were removed as were immunoglobulin genes (likely to be called as differentially expressed due to CLL clonality and variable IG gene usage) (**Fig S2B and S2C**). Differential gene expression analysis were performed in R v.3.6.1[13]. Counts tables for both mRNA and miRNA were analysed for differential expression amongst CLL subgroups using EdgeR v3.32.1[14,15] and custom R code. Batch correction was performed by blocking for batch during multi factor GLM generation for differential expression analysis, or by performing COMBAT correction[16] on normalised expression values.

miRNA:mRNA interaction analysis was performed using the R package miRComb[17] but heavily modified with custom R code and additional miRNA targets databases (including miRTarBase v7.0, IPA expert validation, TargetScan v7.0, miRSVR, miRDB v5.0 and miRRecords[18–22]. Ingenuity pathway analysis (http://www.ingenuity.com/index.html) and DAVID v8[23,24] were used to further interrogate predicted targets. Interaction maps were produced using CytoScape v3[25] to show negatively correlated miRNA:mRNA pairs present in at least one database of miRNA targets. Statistical analysis of miRNA:mRNA interaction counts was performed by quantifying miRNA:mRNA interactions in TargetScan v7.0 and miRDB v5.0 for miRNAs/mRNAs of interest compared to 50,000 cycles of size matched, randomly selected miRNAs/mRNAs using a 1-way student’s t-test. All analyses were performed in R v.3.6.1 using custom code. Data available on request from the authors.
